# The deacylase SIRT5 supports melanoma viability by regulating chromatin dynamics

**DOI:** 10.1101/2020.09.07.286526

**Authors:** William Giblin, Lauren Bringman-Rodenbarger, Angela H. Guo, Surinder Kumar, Alexander C. Monovich, Ahmed M. Mostafa, Mary E. Skinner, Michelle Azar, Ahmed S.A. Mady, Carolina H. Chung, Namrata Kadambi, Keith-Allen Melong, Ho-Joon Lee, Li Zhang, Peter Sajjakulnukit, Sophie Trefely, Erika L. Varner, Sowmya Iyer, Min Wang, James S. Wilmott, H. Peter Soyer, Richard A. Sturm, Antonia L. Pritchard, Aleodor Andea, Richard A. Scolyer, Mitchell S. Stark, David A. Scott, Douglas R. Fullen, Marcus W. Bosenberg, Sriram Chandrasekaran, Zaneta Nikolovska-Coleska, Monique E. Verhaegen, Nathaniel W. Snyder, Miguel N. Rivera, Andrei L. Osterman, Costas A. Lyssiotis, David B. Lombard

**Affiliations:** Department of Pathology, University of Michigan, Ann Arbor, MI 48109 USA; Department of Human Genetics, University of Michigan, Ann Arbor, MI 48109 USA; Department of Biochemistry, Faculty of Pharmacy, Ain Shams University, Cairo, Egypt; Department of Biomedical Engineering, University of Michigan, Ann Arbor, MI 48109 USA; Department of Molecular and Integrative Physiology, University of Michigan, Ann Arbor, MI 48109 USA; Department of Cancer Biology, University of Pennsylvania, Perelman School of Medicine, Philadelphia, PA 19104 USA; Center for Metabolic Disease Research, Department of Microbiology and Immunology, Temple University, Lewis Katz School of Medicine, Philadelphia, PA 19140 USA; Department of Pathology and MGH Cancer Center, Massachusetts General Hospital and Harvard Medical School, Boston, MA 02115 USA; Melanoma Institute Australia, The University of Sydney, Sydney, New South Wales, 2065 Australia; The University of Queensland Diamantina Institute, The University of Queensland, Dermatology Research Centre, Brisbane, QLD 4102 Australia; Department of Dermatology, Princess Alexandra Hospital, Brisbane, Queensland 4102 Australia; Institute of Health Research and Innovation, University of the Highlands and Islands, An Lóchran, 10 Inverness Campus, Inverness, IV2 5NA United Kingdom; Oncogenomics, QIMR Berghofer Medical Research Institute, Brisbane, Queensland, 4006 Australia; Department of Dermatology, University of Michigan, Ann Arbor, MI 48109 USA; Tissue Pathology and Diagnostic Oncology, Royal Prince Alfred Hospital, and NSW Pathology, Sydney, New South Wales, 2050 Australia; Faulty of Medicine and Health, The University of Sydney, Sydney, New South Wales, 2006, Australia; Sanford Burnham Prebys Medical Discovery Institute, La Jolla, California 92037 USA; Departments of Pathology and Dermatology, Yale University School of Medicine, New Haven, CT 06510 USA; Program in Chemical Biology, University of Michigan, Ann Arbor, MI 48109 USA; Center for Computational Medicine and Bioinformatics, University of Michigan, Ann Arbor, MI 48109 USA; Rogel Cancer Center, University of Michigan Medical School, Ann Arbor, MI 48109 USA; Broad Institute of Harvard and MIT, Cambridge, MA 02142 USA; Division of Gastroenterology, Department of Internal Medicine, University of Michigan, Ann Arbor, MI 48109 USA; Institute of Gerontology, University of Michigan, Ann Arbor, MI 48109 USA

**Author notes:** Correspondence: University of Michigan, 3015 BSRB, 109 Zina Pitcher Place, Ann Arbor, MI 48109. These authors contributed equally (Lauren Bringman-Rodenbarger, Angela H. Guo, Surinder Kumar, Alexander C. Monovich, Ahmed M. Mostafa, Mary E. Skinner).

## Abstract

Cutaneous melanoma remains the most lethal skin cancer, and ranks third among all malignancies in terms of years of life lost. Despite the advent of immune checkpoint and targeted therapies, only roughly half of patients with advanced melanoma achieves a durable remission. SIRT5 is a member of the sirtuin family of protein deacylases that regulate metabolism and other biological processes. Germline *Sirt5* deficiency is associated with mild phenotypes in mice. Here we show that SIRT5 is required for proliferation and survival across all cutaneous melanoma genotypes tested, as well as uveal melanoma, a genetically distinct melanoma subtype that arises in the eye and is incurable once metastatic. Likewise, SIRT5 is required for efficient tumor formation by melanoma xenografts and in an autochthonous mouse *Braf;Pten*-driven melanoma model. Via metabolite and transcriptomic analyses, we find that SIRT5 is required to maintain histone acetylation and methylation levels in melanoma cells, thereby promoting proper gene expression. SIRT5-dependent genes notably include *MITF*, a key lineage-specific survival oncogene in melanoma, and the *c-MYC* proto-oncogene. SIRT5 may represent a novel, druggable genotype-independent addiction in melanoma.

## Introduction

Cutaneous melanoma remains the most lethal skin cancer. In 2021, there will be an estimated 106,110 new melanoma cases and 7,180 melanoma-related deaths in the US (1) Melanoma incidence is rising (2), and melanoma ranks third among all cancers in terms of years of life lost (3, 4). Despite the advent of immune checkpoint and targeted therapies, only about half of patients with advanced melanoma achieves long-term remission, even with optimal immune checkpoint therapy (5). Uveal melanoma represents a genetically and clinically distinct subtype of melanoma that arises in the eye, and currently has no effective treatment options once metastatic (6). New therapeutic strategies for advanced melanoma are urgently needed.

Mammalian sirtuins are a family of seven NAD^+^-dependent lysine deacylases that regulate diverse processes to promote cellular and organismal homeostasis and stress responses. Among these proteins, Sirtuin 5 (SIRT5) has remained a somewhat enigmatic and poorly characterized sirtuin. SIRT5 is atypical, in that it lacks robust deacetylase activity, and primarily functions to remove succinyl, malonyl, and glutaryl modifications from lysines on its target proteins, in mitochondria and throughout the cell, thereby regulating multiple metabolic pathways (7–14).

SIRT5-deficient mice are viable, fertile, and mostly healthy (15, 16), with the most prominent effects described to date occurring in the myocardium (17). *Sirt5* knockout (KO) mice are more susceptible to ischemia-reperfusion injury and exhibit impaired recovery of cardiac function compared to wild type (WT) mice (18). Aged *Sirt5* KO mice develop cardiac hypertrophy and mildly impaired ejection fraction (19). Whole-body *Sirt5* KOs, but not cardiomyocyte-specific KOs, show increased lethality in response to cardiac pressure overload (20, 21). Overall, however, the lack of strong phenotypes associated with SIRT5 loss-of-function in normal tissues has hindered progress in understanding the biological significance of SIRT5 and its target post-translational modifications.

Multiple sirtuins are now linked to neoplasia, as tumor suppressors and/or oncogenes, in a context-specific manner (22). In the context of melanoma, genetic inhibition of *SIRT1* in human melanoma cell lines induces senescence and sensitizes drug-resistant cells to vemurafenib, an FDA-approved therapy for the treatment of *BRAF*-mutant melanoma (23). Conversely, genetic *SIRT2* inhibition results in vemurafenib resistance in *BRAF*-mutant melanoma cells by altering MEK/ERK signaling (24). SIRT3 has likewise been reported to play an oncogenic role in melanoma. Reduction of SIRT3 levels in human melanoma lines results in decreased viability, increased senescence and impaired xenograft formation (25). SIRT6 is upregulated in melanoma cells and tissue samples, and SIRT6 depletion in melanoma cell lines results in reduced colony formation and proliferation (26). Paradoxically, *SIRT6* haploinsufficiency induces resistance to targeted therapies in *BRAF*-mutant melanoma cells by regulating IGF/AKT signaling (27).

The functions of SIRT5 in cancer are not well understood, and a subject of active investigation (7). For example, SIRT5 promotes chemoresistance in non-small cell lung carcinoma cells by enhancing NRF2 activity and expression of its targets involved in cellular antioxidant defense (28). SIRT5 promotes Warburg-type metabolism in lung cancer cells by negatively regulating SUN2, a member of the linker of nucleoskeleton and cytoskeleton complex (29). SIRT5 suppresses levels of reactive oxygen species (ROS) via desuccinylation of multiple targets (superoxide dismutase 1, glucose-6-phosphate dehydrogenase, and isocitrate dehydrogenase (IDH) 2), thereby promoting growth of lung cancer cell lines in vitro (30, 31). SIRT5 also plays an important role in facilitating tumor cell growth by desuccinylating serine hydroxymethyltransferase 2 (SHMT2), which catalyzes the reversible, rate-limiting step in serine catabolism, providing methyl groups for cellular methylation reactions via one-carbon metabolism (1CM) (32). Another study indicated that SIRT5 promotes hepatocellular carcinoma (HCC) proliferation and invasion by targeting the transcription factor E2F1 (33). Similarly, it was recently reported that SIRT5 suppresses apoptosis by deacetylation of cytochrome C, thereby promoting HCC growth (34). SIRT5 also promotes breast cancer tumorigenesis by desuccinylating and stabilizing glutaminase (35), an enzyme that catalyzes the conversion of glutamine to glutamate, which supports the metabolic demands of tumorigenesis (36). Another recent publication showed that SIRT5 promotes breast cancer growth in part by suppressing ROS, and described selective SIRT5 inhibitors that markedly impaired tumor growth in vivo (37). In contrast, SIRT5 opposes malignant phenotypes associated with expression of mutant IDH, which generates the novel oncometabolite R-2-hydroxyglutarate, thereby perturbing the epigenome (38). IDH mutant glioma cells show increased protein succinylation, exhibit mitochondrial dysfunction and are resistant to apoptosis. Ectopically expressed SIRT5 in these cells impaired their growth in vitro and in vivo. Another recent report indicates that SIRT5 inactivates STAT3, thus suppressing mitochondrial pyruvate metabolism in lung cancer (39).

Here, we identify a critical requirement for SIRT5 in melanoma cell survival, through chromatin regulation. In all cutaneous and uveal melanoma cell lines tested, from both humans and mice and with varied genetic drivers, SIRT5 depletion resulted in rapid loss of proliferative capacity and cell death. Likewise, SIRT5 loss reduced melanoma formation in xenograft and autochthonous mouse melanoma models. Via transcriptomic analysis, we identified a core set of genes that responds to SIRT5 depletion. Among these, *MITF*, an essential lineage-specific transcription factor in melanoma, is downregulated, along with expression of its targets (40). SIRT5 loss is also associated with reduced expression of *c-MYC*, a well-described proto-oncogene that is often overexpressed in metastatic melanoma and melanoma cell lines, which plays an important role in therapeutic resistance (41, 42). We link the effects of SIRT5 depletion on gene expression to alterations in histone acetylation and methylation induced by metabolic changes occurring in the context of SIRT5 loss-of-function. Taken together, our results identify SIRT5 as a novel genotype-independent dependency in melanoma cells, likely exerting its effects via chromatin modifications and gene regulation. Given the modest effects of SIRT5 loss-of-function in normal tissues, SIRT5 may represent an attractive therapeutic target in melanoma and potentially other cancer types.

## Results

### The chromosomal region encompassing *SIRT5* shows frequent copy number gain in human melanoma

In humans, the *SIRT5* gene localizes to chromosome 6p23. The 6p region exhibits frequent copy number gain in melanoma, an event associated with an adverse prognosis, both in melanoma (43) and other cancers (44). To confirm that gain of the *SIRT5* locus specifically occurs in human melanomas, we mined TCGA (The Cancer Genome Atlas) data (45) using cBioportal, and observed that copy number gain or amplification of *SIRT5* was present in 55% of melanoma cases, whereas *SIRT5* deletion or mutation was rare (Figure 1A, S1A, and S1B). Increased *SIRT5* copy number also correlated with increased *SIRT5* mRNA expression in these samples (Figure S1C). In contrast, the presence of extra copies of the other six sirtuins was much less common in melanoma (Figure 1A and S1A). Activating mutations in *BRAF* and *NRAS* represent the most common oncogenic drivers in cutaneous melanoma (46). *SIRT5* gain or amplification was observed in melanomas with either driver, and in melanomas with the less common driver mutation, *NF1* (Figure 1A). Increased *SIRT5* copy number was associated with moderately worsened overall survival (p=0.0097; Figure 1B), although not progression-free survival (Figure S1D).

**Figure 1.**
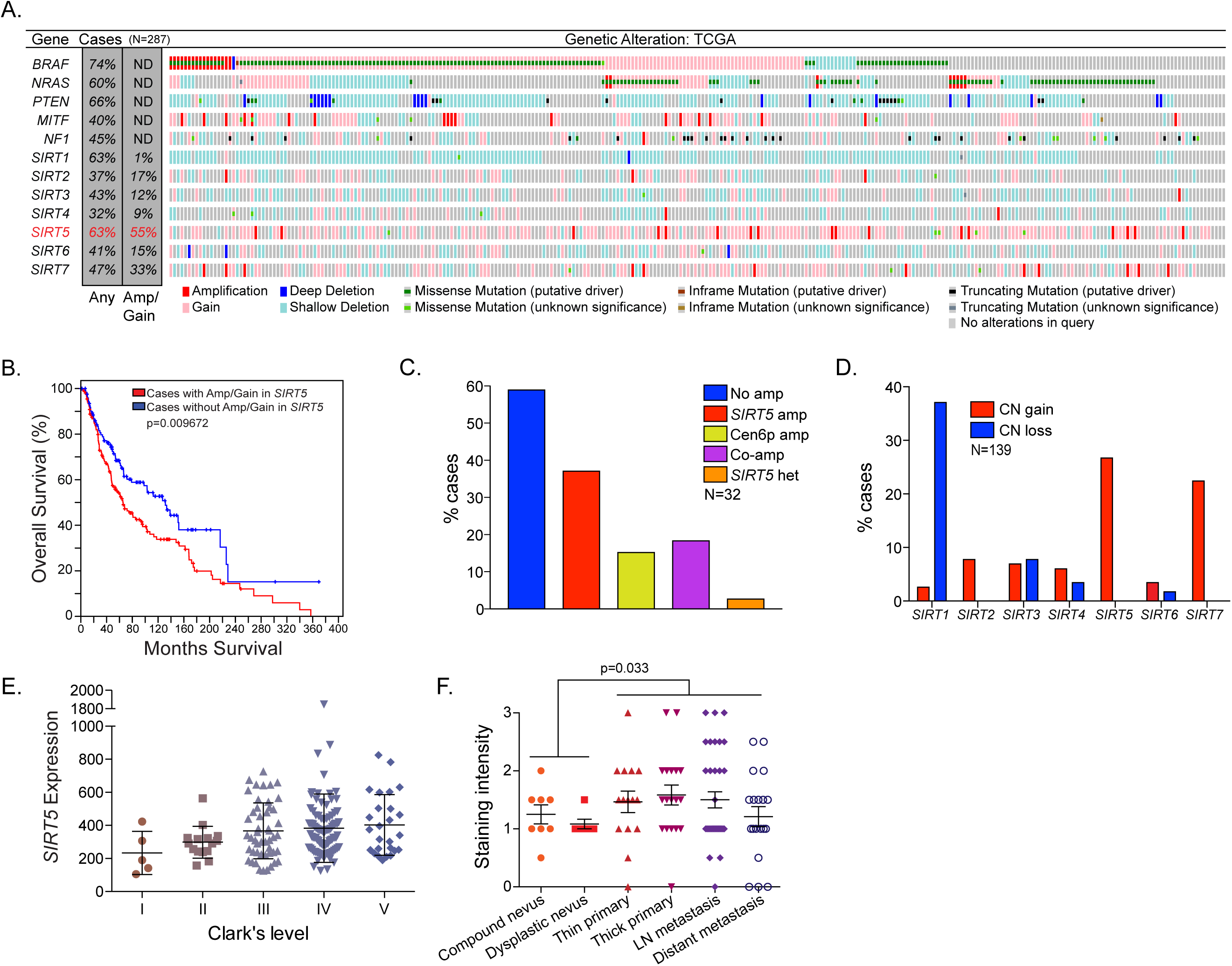
Increased *SIRT5* copy number in human melanoma. **A.** Gain of extra *SIRT5* copies in melanoma. *BRAF*, *NRAS*, *PTEN*, *MITF*, *NF1* and other sirtuins are shown for comparison (n=287; data from TCGA, Provisional, analyzed on cBioPortal). ND, not determined. Percentage of samples with any genomic alteration (Any) or amplification or gain (Amp/Gain) is indicated. Graphed are any alterations queried for the indicated gene. Copy number gain indicates a low-level gain of a single additional copy, and amplification refers to high-level amplification (multiple extra copies). Results from the query (*GENE*: MUT AMP HOMDEL GAIN HETLOSS) in cBioPortal were analyzed and plotted. **B.** Kaplan–Meier analysis of overall survival in melanoma patients with or without copy number gain or amplification of *SIRT5*. Overall survival was analyzed using the query: “*SIRT5*: AMP GAIN.” **C**. *SIRT5* (6p23) and centromere 6p (Cen6p) amplification (amp) or co-amplification (Co-amp) in melanoma, as assayed by FISH staining (n=32). **D.** Sirtuin gene copy number (CN) in human melanoma samples, as assayed by high density SNP array (n=139). **E.** *SIRT5* mRNA expression levels in melanoma correlate with Clark’s level (p=0.0044, linear regression; p=0.037, ANOVA). **F**. SIRT5 protein levels are increased in melanoma relative to benign melanocytic lesions (p=0.0333, Chi-squared; n=14 nevi, n=87 melanoma). See also Figure S1 and Table S1.

To assess further the status of the *SIRT5* locus in melanoma, we performed fluorescence in situ hybridization (FISH) analysis of *SIRT5* and the centromere of chromosome 6 in an independent group of melanoma samples (Figure 1C and S1E). Consistent with TCGA data, increased *SIRT5* copy number was observed in 38% (12/32) of cases analyzed overall, with co-amplification of *SIRT5* and the centromere of chromosome 6 present in 16% (5/32) of cases. Similarly, using comparative genomic hybridization analysis in yet another independent group of melanoma samples, gain of the *SIRT5* locus was present in 27% of melanoma cases analyzed (37/139), the most frequent gain among any of the sirtuins (Figure 1D). We also note that the *SIRT7* locus was amplified in a substantial fraction of melanoma cases. SIRT7 promotes DNA repair by deacetylating and desuccinylating histones (47), however it is not currently known what role SIRT7-mediated deacylation might play in melanomagenesis.

We then interrogated *SIRT5* mRNA expression in melanomas of varied depth of invasion, and found that increased *SIRT5* mRNA expression occurred in melanomas of greater Clark’s level, which are more clinically aggressive and confer a worse prognosis (48, 49) (Figure 1E). Similarly, we examined SIRT5 protein expression in tissue microarrays containing examples of benign and dysplastic nevi, as well as localized and metastatic melanomas. We found by immunohistochemistry that SIRT5 protein was overexpressed in melanomas relative to benign melanocytic lesions (Figure 1F).

To characterize the stage of melanogenesis at which *SIRT5* gain occurs, we screened a panel of genomically characterized benign and dysplastic nevi (n=30) (50) for *SIRT5* somatic mutations and copy number aberrations. No deleterious point mutations were identified in *SIRT5*; however, there was evidence of regional loss of heterozygosity encompassing the *SIRT5* locus in 3/30 benign nevi (10%) assayed (Table S1). However, no *SIRT5* copy number gain or amplification was identified in any of the nevus samples, supporting the idea that *SIRT5* amplification represents a relatively late event in melanomagenesis. This is consistent with the known rarity of such genomic events in nevi (50, 51). Overall, these data show that gain or amplification of *SIRT5* is a common genomic event in melanoma but not nevi.

### SIRT5 is required for survival of BRAF^V600E^ and NRAS^Q61R^ melanoma cells

We assessed the potential requirement of SIRT5 in melanoma cells using a panel of 10 *BRAF or NRAS* mutant melanoma cell lines (Table S2). SIRT5 protein was readily detectable by immunoblot in all cell lines tested (Figure S2A). We initially depleted *SIRT5* using two lentiviral shRNAs targeting distinct regions of the *SIRT5* mRNA (knockdown (KD) 1 and KD2) (11). Although predominantly mitochondrial, SIRT5 is also present in the cytosol and the nucleus (11), and was efficiently depleted from all of these compartments upon *SIRT5* shRNA transduction in all cell lines tested (Figure S2B and S2C). In both BRAF^V600E^ and NRAS^Q61R^ cells, SIRT5 depletion induced rapid loss of proliferation over the course of 7 days (Figure 2A and S2D). Similar results were obtained in an in vitro colony forming assay (Figure 2B). Vemurafenib is a targeted therapy FDA-approved for treatment of BRAF-mutant melanoma. Patients treated with targeted therapies often rapidly relapse with drug-resistant disease (52). SIRT5 inhibition in a vemurafenib-resistant derivative of the melanoma cell line SK-MEL-239, SK-MEL-239**VR**, induced rapid loss of proliferation upon SIRT5 KD, indicating that these vemurafenib-resistant cells retained SIRT5 dependency (Figure 2A and S2E). To complement shRNA-based studies, and to further evaluate the requirement of melanoma cells for SIRT5, we mutated the *SIRT5* locus via CRISPR-Cas9, using four distinct guide RNAs (gRNAs, G1-G4) targeting SIRT5. Consistent with results obtained using shRNA, a dramatic reduction in colony formation was observed in *SIRT5* mutant populations compared to control (Figure 2C). In contrast, SIRT5 KD in several ovarian cancer cell lines did not induce loss of viability as seen in melanoma, indicating that SIRT5 depletion is tolerated in some cancer types (Figure S2F and S2G).

**Figure 2.**
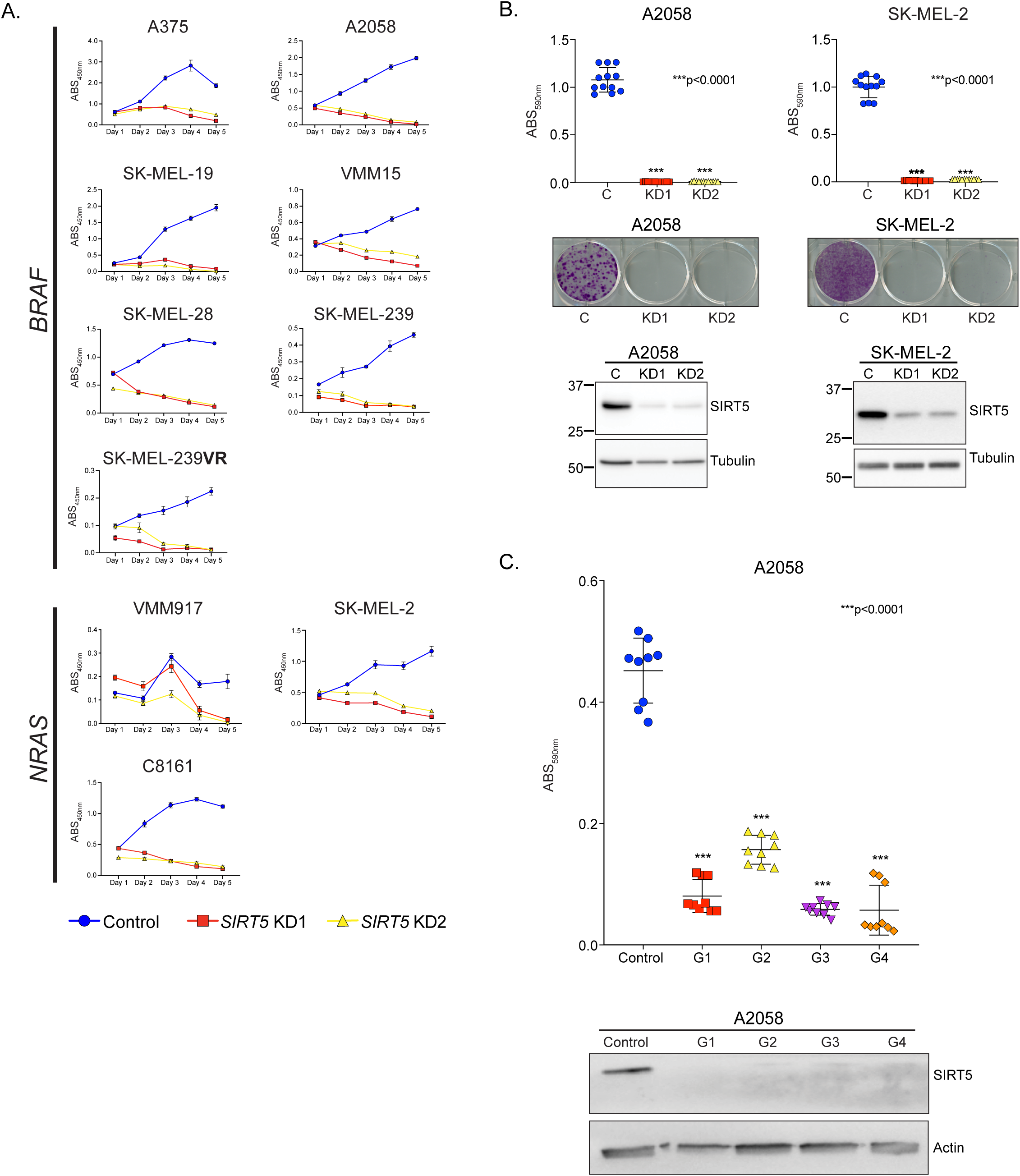
SIRT5 is required for melanoma cell growth and survival. **A.** *BRAF* or *NRAS* mutant melanoma cell lines indicated were infected with a non-targeting shRNA (control) or one of two SIRT5 shRNAs (KD1 or KD2). Equivalent cell numbers were then plated 48 hrs. post-transduction into 96-well plates in the presence of puromycin. Cell mass was determined at the indicated timepoints via WST-1 assay, with absorbance measured at 450nm. Average results (n=6/timepoint) are graphed. Error bars represent standard deviation. Representative of 5/5 SIRT5 shRNAs tested (see also Figure 3B). **B.** SIRT5 KD results in significantly (p<0.0001, unpaired Student’s t-test) impaired colony formation by A2058 and SK-MEL-2 cells 12 days post-transduction. Cell mass was assayed using crystal violet staining, with absorbance measured at 590nm. Average of n=12 technical replicates results are plotted. Error bars represent SD. Representative crystal violet-stained wells are shown. Lower panel, representative immunoblot analysis demonstrating SIRT5 KD. **C.** Top panel, viability of A2058 cells transfected with the indicated CRISPR guide RNA (Control or G1-G4). Cell mass was assayed using crystal violet staining, with absorbance measured at 590nm. Average of n=9 technical replicates results are plotted. Error bars represent standard deviation. Significance calculated using unpaired Student’s t-test. Bottom panel, representative immunoblot analysis confirming CRISPR-mediated SIRT5 loss (Control: empty vector).

### Loss of SIRT5 leads to apoptotic cell death in cutaneous and uveal melanoma cells

We evaluated the mechanism of cellular attrition induced by SIRT5 loss-of-function. SIRT5 depletion in melanoma cells induced cleavage of caspase 3 (Figure 3A and 3B) and induction of Annexin V positivity (Figure 3C and 3D). Importantly, SIRT5 depletion also blocked proliferation and induced cleavage of caspase 3 in uveal melanoma cell lines (Figure 3B; top panel; representative of 4/4 uveal melanoma cell lines tested; see Table S2). Cell loss and induction of caspase 3 cleavage at 96 hrs. post-transduction were also observed, to varying degrees, with an additional 3 unique shRNAs targeting human SIRT5 (Figure 3B; middle panel), and 5 unique shRNAs targeting murine SIRT5 in YUMM5.2, a mouse melanoma cell line (53) (Figure 3B; bottom panel).

**Figure 3.**
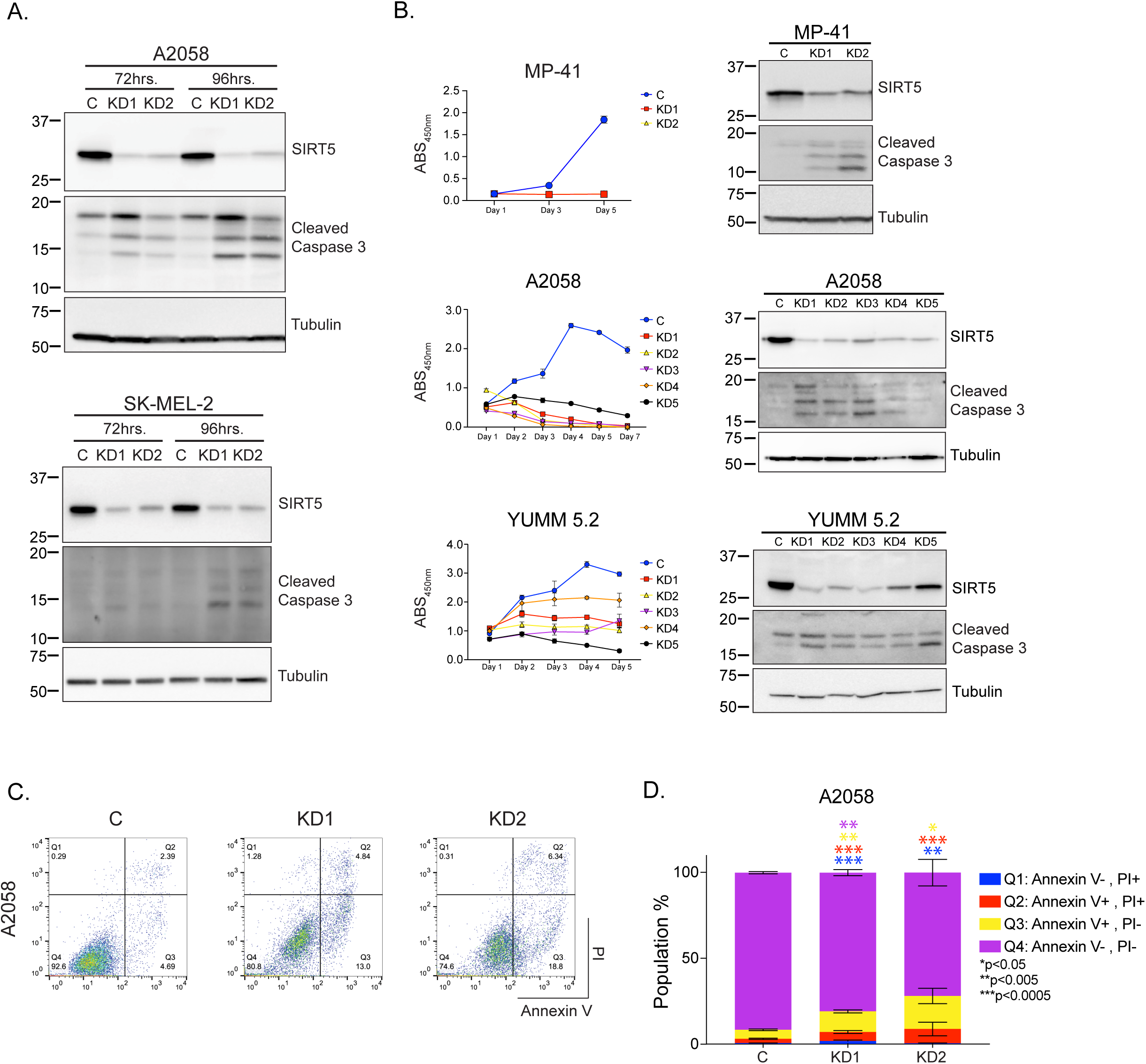
SIRT5 depletion rapidly induces apoptosis in melanoma cells. **A.** Immunoblot analysis demonstrating induction of caspase 3 cleavage 72 and 96 hrs. post-transduction with shRNAs targeting *SIRT5* (KD1 or KD2) in A2058 and SK-MEL-2 cell lines. **B**. Viability of MP-41, A2058 or YUMM5.2 cells infected with control (C) or one of five SIRT5 shRNAs (KD1-KD5) against human *SIRT5* (top and middle panels) or mouse *Sirt5* (bottom panel). Average results (n=6/timepoint) are graphed. Error bars represent standard deviation. Right panels: immunoblot analysis demonstrating loss of SIRT5 and induction of caspase 3 cleavage following SIRT5 KD. **C.** Flow cytometric analysis of A2058 cells stained with Annexin V and propidium iodide (PI), as indicated, showing an increased fraction of Annexin V-positive cells 96 hrs. after SIRT5 KD. **D.** Average of n=3 technical replicates is plotted. Error bars represent (SD). Significance calculated using unpaired Student’s t-test. Increased Annexin V^+^ staining is observed in both the PI-positive and PI-negative populations.

Non-apoptotic mechanisms of cell death have been described in melanomas and other cancer types, specifically: autophagic, ER-stress induced, necroptosis and pyroptosis (54). We evaluated whether SIRT5 KD in melanoma cell lines harboring either BRAF or NRAS mutations induced these alternate cell death pathways. We did not observe increased conversion of LC3 A/B I to LC3 A/B II or SQSTM1/p62 loss (Figure S3A), increased expression of PERK, Calnexin, IRE1 alpha or PDI (Figure S3B), phosphorylation of either MLKL or RIP (Figure S3C), or accumulation of gasdermin D or caspase 1 cleavage products (Figure S3D), although there was some variation observed between different SIRT5 KD constructs. Thus, SIRT5 is required for survival and proliferation of multiple genetically diverse melanoma cell lines in vitro, in both human and mouse, and for survival of human uveal melanoma cells.

### SIRT5 supports robust melanoma tumor formation in vivo

To investigate the potential requirement for SIRT5 to support melanoma tumor development in vivo, we initially employed a xenograft assay. Immediately following transduction with SIRT5 shRNAs, A2058 melanoma cells were subcutaneously injected into the flanks of female NOD/SCID mice (Figure S4A); tumor growth was followed by serial measurement of tumor volume (Figure 4A; left panel). SIRT5 depletion greatly impaired tumor growth and reduced tumor size at endpoint relative to controls (Figure 4A; right panel, 4B and S4B).

**Figure 4.**
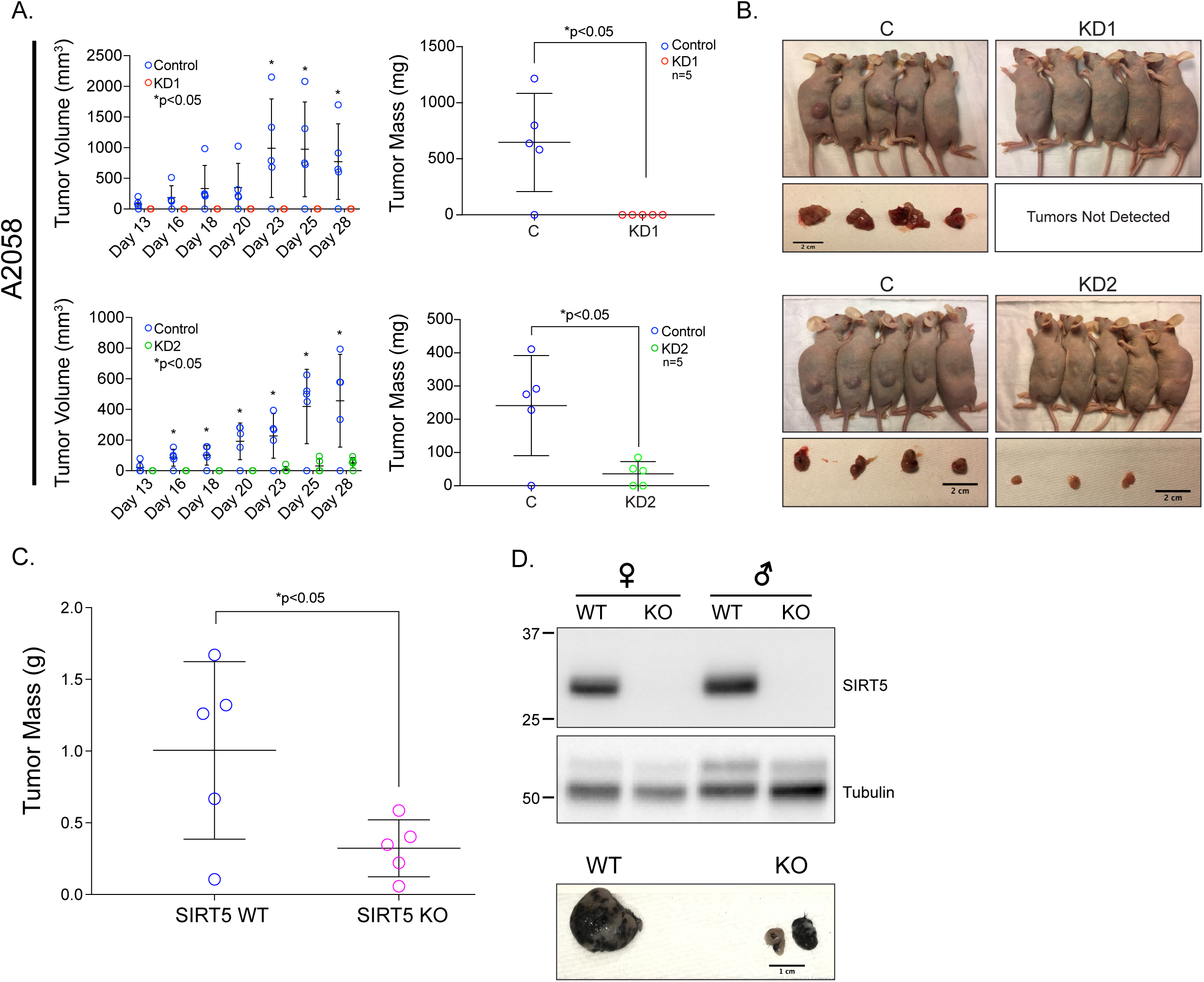
SIRT5 loss-of-function inhibits melanoma tumor growth in vivo. **A.** SIRT5 depletion in A2058 cells results in attenuated xenograft tumor growth. Quantification of tumor size was initiated on day 13 after initial injection of cells (left panel). Tumor size was recorded with Vernier calipers on the days indicated. Each point represents the measurements on n=5 mice for each condition (C, KD1, or KD2). Pairwise representation of endpoint tumor size in each mouse within each group is plotted (right panel). Average tumor mass measurements at day 28 are plotted (p<0.05, paired two-tailed t-test for each group). Error bars represent SD. **B.** Mice were sacrificed, and tumors were dissected at 28 days after initial injection. Scale bar below tumors=2cm. **C**. SIRT5 deficiency attenuates tumor formation in an autochthonous melanoma model. *Sirt5* deficient-mice were bred into the *Braf^CA^;Pten^fl/fl^;Tyr::CreER* background (55). Melanomas were induced in littermate male *Sirt5* WT or *Sirt5* KO mice as shown by topical application of 4HT at ages 4-9 weeks; tumors were weighed following euthanasia. Averages of 5 sets of male mice are plotted (p<0.05, paired two-tailed t-test). Means ± SD are shown. **D.** SIRT5 immunoblot of a representative tumor from a *Sirt5* WT or KO male or female mouse (left panel). Representative tumor from a *Sirt5* WT or KO male mouse, as indicated, after 4HT induction (right panel). Scale bar=1cm.

To examine the role of SIRT5 in melanoma development in a more physiologic, immunocompetent context, we crossed *Sirt5* KO mice to a commonly-used mouse melanoma model, the *Braf^CA^;Pten^fl/fl^;Tyr::CreER* strain (55). Topical application of 4-hydroxytamoxifen (4HT) in this system induces activated BRAF expression and ablation of *Pten* in melanocytes, resulting in melanoma development. In males, SIRT5-deficient mice showed an approximately 3-fold reduction in tumor mass on average (WT: 1.005±0.618g vs. KO: 0.323±0.198g, p<0.05, (Figure 4C and 4D). In our colony, female mice showed rapid ulceration of even small melanoma tumors following induction (not shown), requiring euthanasia of the host and rendering it difficult to assess the effects of SIRT5 in melanoma in females. Thus, SIRT5 promotes human and mouse melanoma growth, both in cell culture and in vivo.

### Neither glucose nor glutamine metabolism are greatly altered by SIRT5 loss

Initially, we considered the possibility that SIRT5 depletion might induce global metabolic collapse and energetic catastrophe in melanoma. SIRT5 has been reported to promote mitochondrial respiration (56, 57) and glycolysis (14). We previously showed that SIRT5 suppresses mitochondrial respiration through Pyruvate Dehydrogenase and Complex II in 293T cells and liver mitochondria (11), a finding recapitulated in some systems (39), but not others (56, 57). We used the XFe96 Extracellular Flux Analyzer to assess the effects of SIRT5 depletion on cellular bioenergetics in melanoma cells. Relative to SIRT5-proficient controls, SIRT5 KD A2058 or A375 cells did not show consistent changes in the extracellular acidification rate (ECAR), a measure of cellular glycolysis (Figure 5A). Likewise, glucose-dependent mitochondrial oxygen consumption rate (OCR), ATP production, and mitochondrial membrane potential were not consistently affected by SIRT5 depletion (Figure 5B-5D).

**Figure 5.**
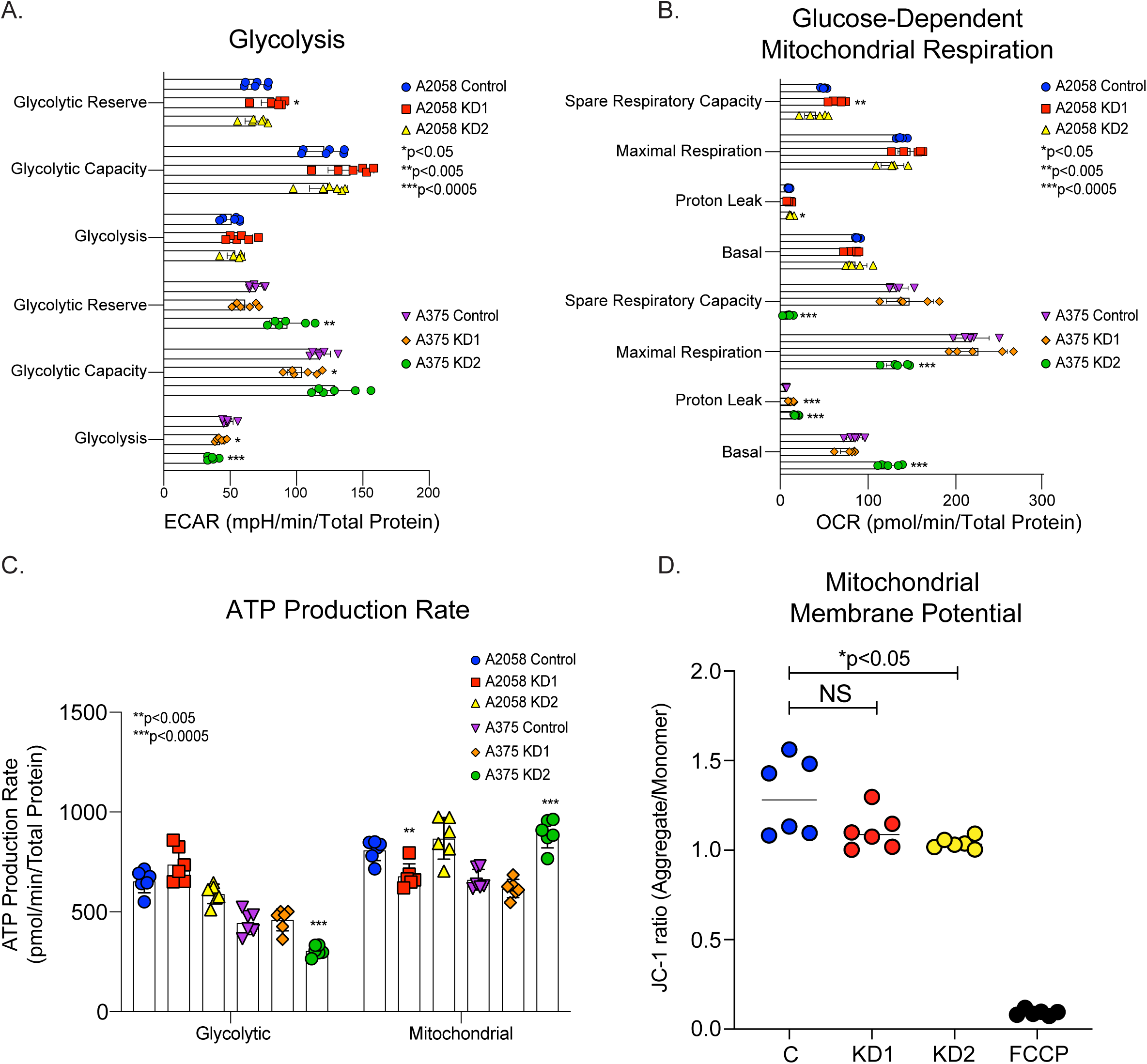
Bioenergetics are maintained upon SIRT5 loss in melanoma cells. A2058 and A375 cells maintain glycolytic function (**A.**), glucose-dependent mitochondrial respiration (**B**.), and ATP production (**C.**) upon SIRT5 depletion compared to control cells. Mitochondrial respiration, glycolytic stress tests, and ATP production rates were measured at 72 hrs. post-transduction with shRNAs against SIRT5 using a Seahorse XFe96 Analyzer. All rates are normalized to total protein content per sample (n=6 for **A.** and **C.**, n=5 for **B.**). OCR, oxygen consumption rate; ECAR, extracellular acidification rate. Error bars represent standard deviation. **D.** Mitochondrial membrane potential is stable in A2058 cells after SIRT5 loss (C, control cells, n=6). Cells were incubated with JC-1, a dye which exhibits membrane potential-dependent accumulation in mitochondria, indicated by a fluorescence emission shift from green to red. Mitochondrial depolarization is indicated by a decrease in the red:green (aggregate:monomer) fluorescence intensity ratio. FCCP, a mitochondrial uncoupler, depolarizes mitochondrial membrane potential and is used as a positive control. Significance calculated using unpaired Student’s t-test.

Melanoma and many other cancer types replenish the TCA cycle in part via glutaminolysis (58–61). In this pathway, glutaminase (GLS) catalyzes conversion of glutamine to glutamate, generating carbon and nitrogen to fuel the metabolic demands of tumorigenesis. In breast cancer cells, SIRT5 desuccinylates GLS to stabilize it, protecting it from ubiquitination and subsequent degradation. Loss of SIRT5 resulted in decreased GLS expression, exogenous glutamine consumption, glutamine-derived intracellular metabolite levels, and cellular proliferation (35). These findings, along with reports that inhibiting glycolysis or glutamine metabolism sensitizes melanoma cells to cell death, prompted us to investigate a potential role for SIRT5 in promoting glutamine metabolism in melanoma (62, 63). We cultured control or SIRT5 KD A2058 cells in medium containing glutamine labeled with stable isotopes ([^15^N_2_]-glutamine or [^13^C_5_]-glutamine) or in medium containing [^13^C_6_]-glucose, and measured both labeling derived from ^13^C or ^15^N and total quantities of cellular metabolites. Following SIRT5 KD, the fractional labeling of glutamine derived metabolites (glutamate, aspartate and TCA cycle metabolites) modestly decreased when cells were cultured in [^13^C_5_]-glutamine, but showed corresponding (or compensatory) increases in labeling when cultured in ^13^C_6_-glucose (Figure S5A-S5B). Labeling derived from ^15^N_2_-glutamine was inconsistent between the two KD constructs analyzed in A2058 cells (Figure S5C). Overall, these results are consistent with previous results showing that SIRT5 promotes glutaminase activity (35). However, importantly, total cellular amounts of glutamate, aspartate, TCA cycle or other metabolites were not consistently reduced by SIRT5 loss (Figure S5D), indicating that although the depletion of SIRT5 may reduce glutaminase activity, this effect is insufficient to compromise levels of essential cellular metabolites. In parallel studies, glutamine-dependent mitochondrial OCR, and GLS protein levels, were assessed, and were not appreciably altered by SIRT5 depletion across multiple melanoma cell lines (Figure S5E and S5F). Moreover, incubation of SIRT5 KD A2058 cells with exogenous non-essential amino acids plus alpha-ketoglutarate – interventions that can rescue defects in glutamine catabolism (61) – in the context of SIRT5 KD failed to rescue the proliferative defect observed upon SIRT5 loss (Figure S5G). Taken together, these data indicate that neither glycolysis nor glutamine metabolism represent major SIRT5 target pathways in promoting melanoma viability.

### Transcriptomic analysis reveals a requirement for SIRT5 in supporting *MITF* and *MITF* target gene expression

To understand the requirement of melanoma cells for SIRT5 mechanistically, RNA-seq based transcriptomic analysis was performed on three cutaneous melanoma cell lines (A2058, A375 and SK-MEL-2), each subjected to SIRT5 depletion using two distinct shRNAs (Table S3). A gene was scored as differentially expressed only if it was consistently altered in all biological replicates by both independent SIRT5 shRNAs. We identified core sets of protein-coding genes whose expression responded to SIRT5 KD, many of which overlapped among the cell lines (Figure 6A and 6B). We then asked if any significant differentially expressed genes (DEGs) found in our SIRT5 RNA-seq dataset correlated with *SIRT5* expression in TCGA data of clinical human skin cutaneous melanoma samples (see Methods for details). The most significant positively correlated overlapping DEG in this analysis was the Melanocyte Inducing Transcription Factor (*MITF*) (Figure 6C).

**Figure 6.**
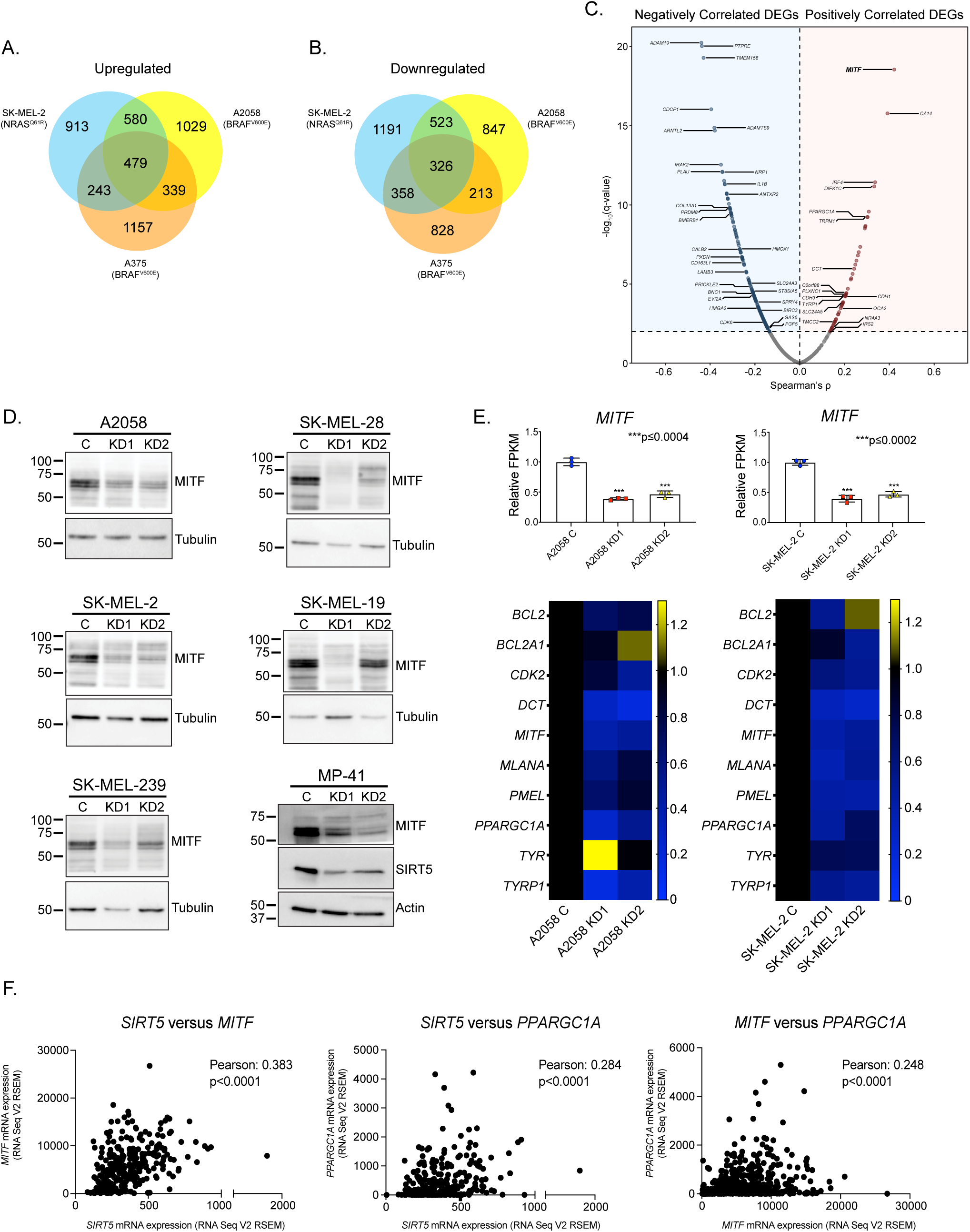
Expression of MITF and MITF target genes is dependent upon SIRT5. Genes (**A.**) upregulated or (**B.**) downregulated upon SIRT5 KD. Only genes significantly (p<0.05) altered in both KDs in each cell line, as indicated, were scored. **C.** Expression levels of differentially expressed genes (DEGs; qadj<0.05) in response to SIRT5 KD were correlated with *SIRT5* gene expression using Spearman’s rank correlation coefficient in 443 sequenced human skin cutaneous melanoma (SKCM) samples, identifying DEGs with significant clinical correlation with SIRT5 expression (q<0.01). Labeled genes represent oncogenes or extremely correlated genes most significantly altered by SIRT5 knockdown (q<0.0001, log2FoldChange>2). **D.** Immunoblot demonstrating loss of MITF expression 96 hrs. post-transduction with shRNAs targeting *SIRT5* (KD1 or KD2) compared to a non-targeting control (C) in 5 cutaneous and one uveal melanoma cell lines, as indicated. **E.** Relative FPKMs in A2058 and SK-MEL-2 cells demonstrate a loss of *MITF* (bar graphs, upper panels) and several *MITF* target gene transcripts upon SIRT5 KD (heatmaps, lower panels). Scale bars adjacent to heat maps indicate linear fold change (control (C) set to 1). Significance calculated using unpaired Student’s t-test. **F**. Expression of *SIRT5*, *MITF* and the *MITF* target, *PPARGC1A* are positively correlated in melanoma clinical samples (p<0.0001, data from TCGA, analyzed on cBioPortal; see Figure 1A).

*MITF,* a key lineage-specific oncogenic transcription factor in melanoma that plays crucial roles in development and proliferation of melanocytes (40). MITF is expressed in human melanomas, and *MITF* amplification, present in a subset of melanoma tumors, portends a poor prognosis (64). Melanomas exhibit a wide range of MITF expression levels (65–67). In cutaneous melanoma cells with robust baseline MITF expression, MITF protein and mRNA expression declined markedly in response to SIRT5 KD (Figure 6D and 6E), associated with decreased expression of MITF’s canonical targets: genes involved in metabolism (*PPARGC1A*), melanocytic differentiation (*TYR*, *MLANA*), cell survival (*BCL2*) and others (Figure 6E). A trend towards a reduced *MITF* gene expression profile was also observed in A375 cells, which have low baseline MITF expression, upon SIRT5 KD (Figure S6A). Decreased *SIRT5*, *MITF* and *MITF* target gene expression was validated by qRT-PCR in A2058 cells, and to a lesser degree, in SK-MEL-2 (Figure S6B). We also observed a decrease in MITF protein levels upon SIRT5 KD in MP-41 cells, a uveal melanoma line (Figure 6D).

To assess the potential relationship between SIRT5 and MITF in a more physiologic, non-loss-of-function setting, we mined TCGA data, to test whether any correlation exists between *SIRT5* and *MITF* mRNA expression in melanoma clinical samples. Consistent with the RNA-seq data, mRNA co-expression analysis revealed a strong positive correlation between *SIRT5*, *MITF* and two canonical *MITF* target genes, *PPARGC1A* and *BCL2*. Indeed, the correlation between *SIRT5* and *MITF* expression was stronger than that of *MITF* with these two of MITF’s targets (Figure 6F and S6C). As a specificity control, *SIRT3* levels showed a modest, negative correlation with *MITF* expression (Figure S6C). These data suggest that SIRT5 expression levels influence expression of *MITF* and its targets in patient melanoma tumors.

Previous reports demonstrate that the proto-oncogene *c-MYC* is upregulated in melanoma tumors and cell lines, acting to bypass mutant BRAF-or NRAS-induced senescence during melanomagenesis (41). Furthermore, siRNA KD of c-MYC in melanocytic tumor cells results in a loss of MITF expression (68). Consistent with these data, we observed a loss of MITF expression and a concomitant reduction in expression of *c-MYC* in SIRT5-depleted melanoma cell lines. A positive correlation between *SIRT5* and *c-MYC* RNA expression in melanoma tumors from TCGA data was observed (Figure S6C). Both c-*MYC* RNA and c-MYC protein levels were decreased in melanoma cells after SIRT5 ablation (Figure S6D and S6E).

Gene set enrichment analysis (GSEA) was used to identify pathways affected by SIRT5 depletion. GSEA revealed negatively enriched gene patterns in *c-MYC*, c-MYC-target gene signatures, and mitochondrial biogenesis pathways (Figure S6F). We also observed a positive enrichment of genes involved in apoptosis, consistent with our observation that SIRT5 loss induces apoptosis in melanoma cells (see Figure 3). Ingenuity Pathway Analysis (IPA) of transcriptional regulators predicts that both *MITF* and *c-MYC* were significantly inhibited by SIRT5 depletion, based on comparisons between data from aggregated *SIRT5* KD melanoma cells and *SIRT5* control lines (Figure S6G). The multiple canonical pathways altered upon SIRT5 loss highlight other, pleiotropic effects of SIRT5 depletion on melanoma cells (Figure S6H). Taken together, these data show that SIRT5 promotes expression and activity of two key oncogenic drivers, MITF and c-MYC, in melanoma.

### SIRT5 regulates melanoma cell metabolism to promote histone acetylation

To obtain systems-level insight into potential roles for SIRT5 in regulating gene expression, we re-analyzed our transcriptomic data, using a genome-scale model of human metabolism to identify metabolic reactions that change in activity after SIRT5 KD. The Recon1 human network model used contains a relationship between 3,744 reactions, 2,766 metabolites,1,496 metabolic genes, and 2,004 metabolic enzymes (69). This network model has been used successfully to predict the metabolic behavior of various cancer cells and stem cells (70, 71). Using this model, we identified a metabolic flux state most consistent with expression data for each of the three cell lines after SIRT5 depletion. This was achieved by maximizing the activity of reactions that are associated with up-regulated genes and minimizing flux through reactions that are down-regulated for each condition, while simultaneously satisfying the stoichiometric and thermodynamic constraints embedded in the model (see Methods).

The model identified 20 reactions among the 3744 that showed significantly different activity across all cell lines after SIRT5 KD (p<0.01; Figure 7A, Table S4). Among these, the enzyme ATP-Citrate Lyase (ACLY) was predicted to have the most significant change, with reduced activity after SIRT5 KD. ACLY generates acetyl-CoA from citrate, thereby playing an important role in supporting histone acetylation (72). Furthermore, the mitochondrial methylenetetrahydrofolate dehydrogenase reaction was also predicted to have reduced activity after SIRT5 loss, a part of the folate and one-carbon metabolism (1CM) pathways (see below). Several reactions involving cholesterol metabolism and nucleotide salvage were also affected by SIRT5 KD, highlighting the pervasive effects of SIRT5 in melanoma cells.

**Figure 7.**
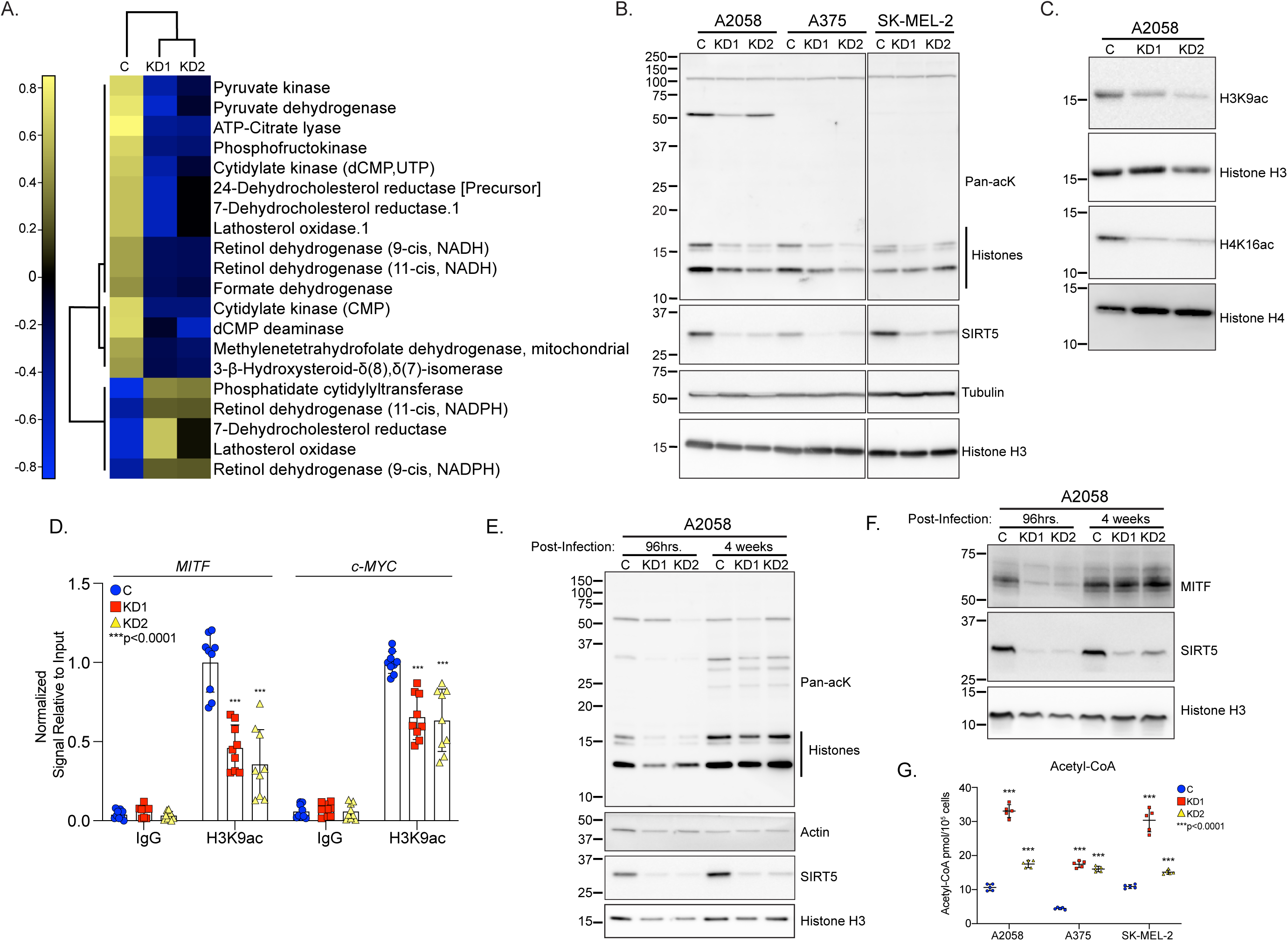
SIRT5 promotes histone acetylation in melanoma. **A.** Heatmap of z-scores calculated from metabolic reaction fluxes predicted by genome-scale modeling to be differentially active (p<0.01) after SIRT5 KD. **B.** Total histone acetylation is reduced 96 hrs. post-transduction with shRNAs targeting *SIRT5* (KD1 or KD2) compared to a non-targeting control (C) in melanoma cell lines. Lanes were run on the same gel but are noncontiguous. **C.** Immunoblot demonstrating loss of H3K9ac and H4K16ac 96 hrs. post-transduction with shRNAs targeting *SIRT5* (KD1 or KD2) compared to a non-targeting control (C) in A2058 cells. **D.** H3K9ac is reduced within the promoter regions of *MITF* and *c-Myc* in SIRT5-depleted A2058 cells via CUT&RUN followed by qRT-PCR. Signal (Ct values) relative to input DNA were normalized to control (C) samples for each primer set. Graphed are averages of n=9 replicates. Error bars represent SD. Significance calculated using unpaired Student’s t-test. Acetylation (**E.**) and MITF expression (**F.**) are restored in A2058 cells lacking SIRT5 after 4 weeks of continual culture in puromycin. **G.** Total cellular acetyl-CoA levels are increased in A2058, A375 and SK-MEL-2 cells 96 hrs. after SIRT5 depletion. Acetyl-CoA abundance was quantified by liquid chromatography-high resolution mass spectrometry and normalized to cell number. Plotted are average (n=5) acetyl-CoA levels as pmol acetyl-CoA/10^5^ cells. Error bars represent SD. Significance calculated using unpaired Student’s t-test.

To test the predictions of the metabolic model, we evaluated protein acetylation levels in SIRT5 KD cells. Indeed, SIRT5 depletion induced a striking decrease in total lysine acetylation, most notably on histones, including H3K9 acetylation (H3K9ac) and H4K16 acetylation (H4K16ac) (Figure 7B and 7C). This reduction in H3K9ac, a known mark of active gene expression (73), combined with the decrease in MITF and c-MYC, prompted us to test whether H3K9ac levels are reduced within the promoter regions of these genes. CUT&RUN (Cleavage Under Targets and Release Using Nuclease) followed by qRT-PCR in A2058 cells demonstrated that upon SIRT5 depletion, a significant reduction of H3K9ac in the promoter regions of both *MITF* and *c-MYC* occurred (Figure 7D), suggesting a role for SIRT5 in maintaining transcriptional activity of these genes in melanoma cells by promoting histone acetylation.

After 4 weeks in culture following SIRT5 KD, a small residual population of A2058 cells overcame SIRT5 loss-of-function to survive and proliferate, although SIRT5 depletion was maintained. Importantly, total lysine acetylation and MITF expression was restored in surviving SIRT5 KD A2058 cell populations (Figure 7E and 7F), consistent with the relevance of SIRT5-driven histone acetylation in melanoma survival. This phenotype was recapitulated in vivo. Although tumors that formed in SIRT5-deficient *Braf^CA^;Pten^fl/fl^;Tyr::CreER* mice were smaller than WT controls (see Figure 4), total lysine acetylation, H3K9ac, MITF and c-MYC protein levels were similar to controls. Markers for cell death (PARP cleavage) and cellular proliferation (PCNA and phospho-histone H3 S10 (H3pS10)) were also similar between SIRT5 WT and SIRT5-deficient tumors in the model, suggesting that these parameters may have recovered during successful tumor formation (Figure S7D).

Protein acetyltransferases employ acetyl-CoA to acetylate their protein targets, including histones (74). To investigate the potential basis for reduced histone acetylation in SIRT5-depleted melanoma cells, we employed a sensitive mass spectrometry-based method to assess total cellular acetyl-CoA levels (75, 76). Surprisingly, we observed an *increase* of total cellular acetyl-CoA after SIRT5 KD (Figure 7G), implying that reduced acetyl-CoA levels do not contribute to the observed decrease in lysine acetylation upon SIRT5 depletion, and suggesting that other phenomena, such as reduced acetyltransferase activity may underlie the reduced acetylation levels in SIRT5-depleted melanoma cells (see Discussion).

### SIRT5 promotes one-carbon metabolism and histone methylation in melanoma

To investigate further how SIRT5 may function to affect gene expression in melanoma, SIRT5-depleted melanoma cell lines were profiled using liquid chromatography coupled tandem mass spectrometry (LC-MS/MS)-based metabolomics, followed by functional analysis using MetaboAnalyst pathway enrichment (Table S5). Two *BRAF* mutant lines (A2058 and A375) and an *NRAS* mutant (SK-MEL-2) showed perturbations in pathways involving 1CM in response to SIRT5 depletion (Figure 8A, S7A and S7B). 1CM is comprised of the linked folate and methionine cycles (77). Outputs include metabolites required for amino acid and nucleotide synthesis; glutathione for antioxidant defense; and crucially, S-adenosylmethionine (SAM) for methylation reactions, including those on histones. We observed a reduction in levels of several key 1CM metabolites upon SIRT5 depletion in *BRAF* mutant melanoma cell lines, but not in SK-MEL-2 (Figure 8B).

**Figure 8.**
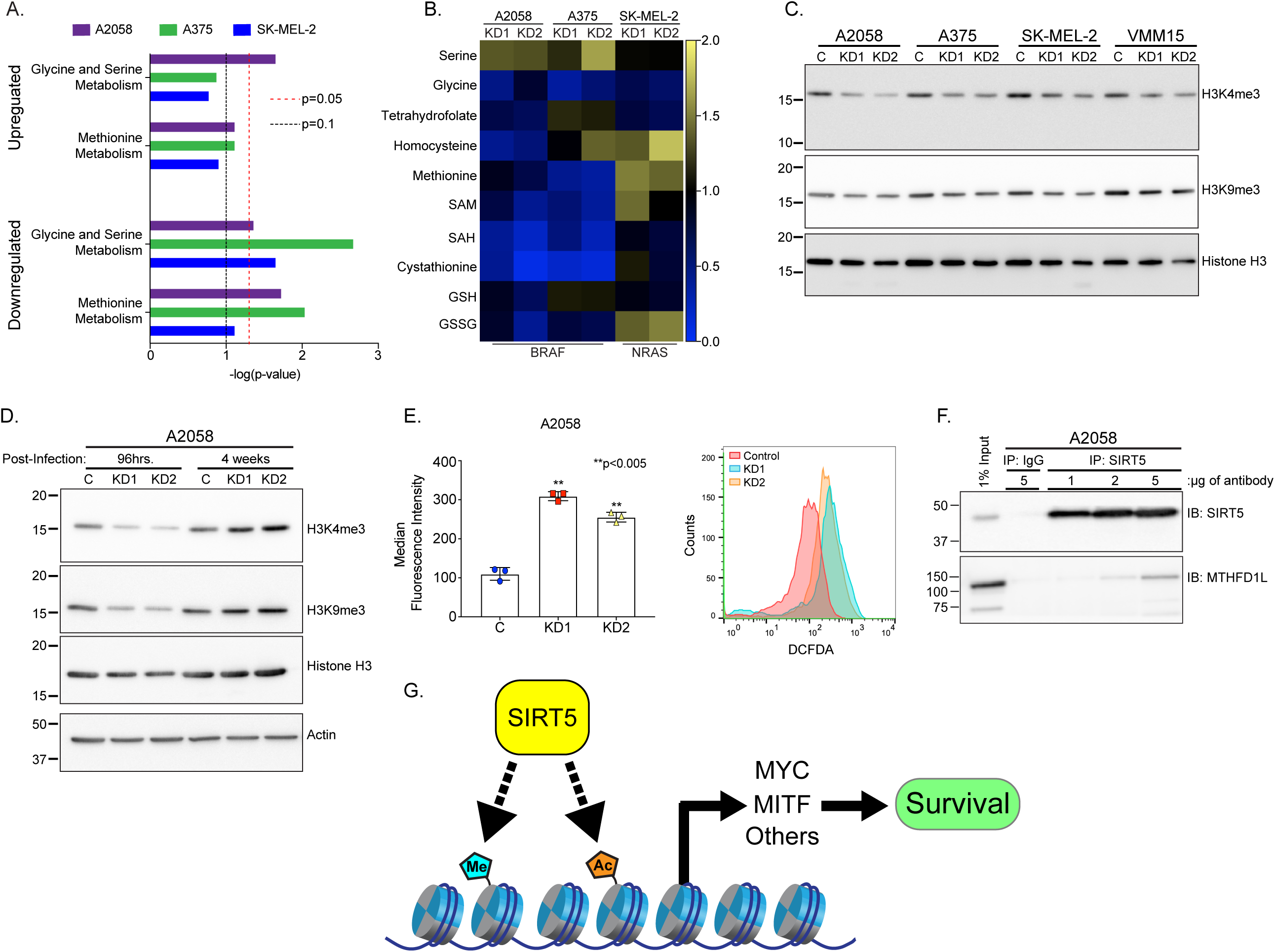
SIRT5 promotes histone methylation and reduced cellular ROS levels in melanoma. **A.** LC-MS/MS-based metabolite profiling followed by MetaboAnalyst pathway analysis demonstrate alterations in glycine and serine and methionine biosynthesis pathways in melanoma cells upon SIRT5 depletion. **B.** Perturbations in 1C metabolite levels in response to SIRT5 loss in the cell lines shown. Each column represents the mean of 3 independently prepared biological replicates. Metabolite levels in SIRT5 depleted (KD1 and KD2, as indicated) samples are normalized to control. SAM, S-adenosyl-methionine; SAH, S-adenosylhomocysteine; GSH, reduced glutathione; GSSG, glutathione disulfide. **C.** H3K4me3 and H3K9me3 immunoblot in melanoma cells 96 hrs. post-transduction with shRNAs targeting *SIRT5* (KD1 or KD2) compared to a non-targeting control (C). **D.** H3K4me3 and H3K9me3 levels are restored in A2058 cells lacking SIRT5 after 4 weeks of continual culture in puromycin. **E.** Flow cytometric analysis of DCFDA-stained A2058 cells 96 hrs. post-transduction with shRNAs targeting *SIRT5* (KD1 or KD2) reveals increased ROS compared to a non-targeting control (C), p<0.005. Left panel, average mean fluorescence intensity of DCFDA positive populations in n=3 samples. Error bars represent SD. Significance calculated using unpaired Student’s t-test. Right panel, representative flow cytometric of A2058 cells stained with DCFDA. **F.** SIRT5 interacts with MTHFD1L in A2058 cells. Increasing amounts of anti-SIRT5 antibody increases SIRT5-MTHFD1L coprecipitation compared to normal rabbit IgG control. Basal expression of SIRT5 and MTHFD1L in whole-cell extract (1% of initial amount used for immunoprecipitation) is shown for comparison. **G.** Proposed model of promotion of MITF and c-MYC expression via SIRT5-dependent chromatin modifications in human melanoma. Me, methylation; Ac, acetylation.

Histone methylation, particularly H3K4 trimethylation (H3K4me3), is highly sensitive to fluctuations in SAM levels (78). We observed reductions in H3K4me3 and H3K9me3 in melanoma cells following SIRT5 KD, consistent with 1CM perturbation (Figure 8C). However, addition of exogenous SAM did not consistently restore H3K4me3 or H3K9me3, nor did it markedly elevate levels of these marks in control cells (Figure S7C and not shown). As for acetylation, SIRT5-depleted melanoma cells that grew out after prolonged culture recovered H3K4me3 and H3K9me3 levels (Figure 8D), while maintaining reduced SIRT5 expression (Figure 7E), suggesting that loss of these histone modifications represents an important driver of the lethality associated with SIRT5 depletion in melanoma.

A decrease in cellular glutathione content occurring in the context of impaired 1CM would be predicted to elevate levels of cellular reactive oxygen species (ROS) (79). Consistently, in A2058 cells, we observed increased staining with 2′,7′-dichlorofluorescin diacetate (DCFDA), a ROS-sensitive dye, following SIRT5 depletion (Figure 8E). However, treatment with the antioxidants, N-acetylcysteine, mitoTEMPOL or β-mercaptoethanol failed to mitigate cell lethality after SIRT5 loss (data not shown), indicating that regulation of ROS levels is not likely a primary determinant of the requirement of melanoma cells for SIRT5.

We noted that previous proteomic surveys identified the 1CM enzyme, MTHFD1L (methylenetetrahydrofolate dehydrogenase (NADP^+^ dependent) 1 like), as a candidate SIRT5 substrate (11, 80). MTHFD1L is a 1CM enzyme that participates in the folate cycle to convert formate and tetrahydrofolate into 10-formyl-tetrahydrofolate in an ATP-dependent reaction. We tested the interaction of MTHFD1L with SIRT5 in the context of melanoma, and found that MTHFD1L co-immunoprecipitates with SIRT5 (Figure 8F). These data suggest a potential role for SIRT5 in regulating multiple 1CM enzymes, such as SHMT2 and potentially MTHFD1L and others, to promote 1CM and histone methylation. Likewise, since SK-MEL-2 cells showed a reduction in histone H3K4me3 levels without apparent declines in 1C metabolites under our experimental conditions, it is likely that SIRT5 plays additional roles in regulating histone methylation, perhaps in an oncogenic driver-dependent manner. We propose that SIRT5 regulates histone methylation and acetylation via regulation of multiple protein targets in melanoma cells.

## Discussion

Sirtuin-family NAD^+^-dependent protein deacylases regulate metabolism and other diverse aspects of cell biology (81). SIRT5 is a poorly-understood, atypical sirtuin, whose primary known biochemical function is to remove succinyl, malonyl, and glutaryl groups from lysines on its target proteins (8, 9, 11–13). A substantial fraction of SIRT5 is present in the mitochondrial matrix; however, SIRT5 is present and functional in the cytosol, and even in the nucleus (11, 14). Most of the phenotypes associated with SIRT5 loss-of-function in normal cells and tissues reported in the literature to date are remarkably mild (17). In sharp contrast, here we report that cutaneous and uveal melanoma cells show exquisite dependency on SIRT5, in a genotype-independent manner. SIRT5 depletion, induced by shRNA or CRISPR/Cas9, provokes dramatic, rapid loss of cell viability and induction of apoptosis in both cutaneous and uveal melanoma cell lines. Likewise, SIRT5 promotes melanoma xenograft tumor formation in immunocompromised mice, and melanoma formation in an autochthonous *Braf*;*Pten*-driven mouse melanoma strain.

Our transcriptomic analyses reveal that SIRT5 plays a major role in maintaining proper gene expression in melanoma cells. SIRT5-dependent genes notably include the lineage-specific oncogenic transcription factor *MITF* (82) and *c-MYC* (41). In the TCGA dataset, *SIRT5* levels correlate with those of *MITF* and *c-MYC*, suggesting that SIRT5 activity influences both *MITF* and *c-MYC* expression in a physiologic context. Indeed, we found that SIRT5 depletion results in loss of H3K9ac, a marker for active transcription, within the promoter regions of these genes. These data are consistent with previously published results describing a role for histone modifications in sustaining MITF expression and melanoma proliferation (83). Genetic or pharmaceutical inhibition of the p300 acetyltransferase results in reduced MITF expression, reduced histone acetylation within of the MITF promoter, and induction of markers of cellular senescence in melanoma cell lines, suggesting regulation of chromatin dynamics as a mechanism of MITF expression and melanoma growth (83). Via metabolomic analysis, we identified a role for SIRT5 in promoting 1CM in two BRAF-dependent cell lines, and in maintaining histone trimethylation at H3K4 and H3K9, marks associated with transcriptional activation and repression, respectively. SIRT5 also plays a distinct role in maintaining histone acetylation. To our knowledge, SIRT5 is the first protein implicated in maintaining both histone methylation and acetylation, highlighting its important roles in maintaining chromatin structure and gene expression in melanoma.

Our in vivo findings in an autochthonous system are in contrast to a published study by Moon *et al.,* in which SIRT5 deficiency was found to exert no impact on tumor growth in a similar mouse melanoma model as the one used in our studies (84). Several potential explanations exist for this discrepancy. Moon *et al.* used a *Sirt5* allele distinct from the one employed in our work; the *Sirt5* allele used in their analysis deletes a single exon in the *Sirt5* gene (16), whereas the one used herein deletes essentially the entire *Sirt5* protein coding sequence (15). Likewise, subtle genetic background differences in the strains of the mice used may contribute to these discrepancies, as could microbiome differences between the mouse colonies. Another potential explanation involves the protocol used to induce gene recombination; we applied a higher concentration of tamoxifen than did Moon *et al.* (64.5 mM versus 5 mM). Importantly, since this model is a global *Sirt5* KO, we cannot rule out the possibility that SIRT5 may function melanoma cell non-autonomously in this system, for example, by modulating the anti-melanoma immune response or other aspects of the tumor microenvironment. However given the striking dependency of cultured melanoma cells on SIRT5 in vitro, we strongly suspect that a very important component of SIRT5’s function, at minimum, is a cell-autonomous pro-survival role in melanoma cells.

MITF is a member of microphthalmia family of transcription factors, and is dysregulated in melanoma (85). Attenuation of melanocyte differentiation and pigmentation are observed in humans and mice deficient for MITF activity, highlighting the importance of MITF in melanocyte survival and function. Likewise, MITF is known to play key roles in melanoma cell survival and differentiation, and *MITF* amplification occurs in 15% to 20% of melanomas, associated with a worsened prognosis (64). In melanoma cell lines where MITF is expressed, SIRT5 depletion induced a rapid decrease in expression of MITF itself and several well-characterized MITF targets. Likewise, in TCGA data, *SIRT5* and *MITF* levels were highly correlated, suggesting that SIRT5 may play a role in regulating MITF in tumors in vivo. Notably, we were unable to rescue the lethality of SIRT5 depletion by overexpressing MITF in melanoma cells (data not shown); however, this experiment is complicated by the fact that MITF overexpression itself can drive melanoma cells to leave the cell cycle and differentiate, and thus is likely selected against in short-term culture (86). Likewise, we were unable to rescue SIRT5-depleted melanoma cells via c-MYC overexpression, although we were able to overexpress c-MYC (data not shown). Nevertheless, given the well-known importance of these transcription factors in melanoma pathobiology, we hypothesize that loss of MITF and c-MYC expression likely represent important mechanisms through which SIRT5 promotes melanoma viability.

We did not observe major effects of SIRT5 depletion on OCR, ECAR, or overall ATP production in melanoma. Instead, through mass spectrometry-based metabolite profiling, we identified one-carbon metabolism (1CM) as one SIRT5 target pathway likely important for maintenance of gene expression and melanoma viability. 1CM consists of the linked folate and methionine cycles. A major output of 1CM is SAM, the universal methyl donor in mammalian cells. Metabolite profiling in two *BRAF* mutant melanoma cells lacking SIRT5 reveals profound perturbations in levels of many 1C metabolites, including reductions in cellular SAM. Moreover, H3K4me3, a mark of active gene expression and a sensitive marker for intracellular SAM levels, drops in response to SIRT5 loss-of-function. Furthermore, global lysine acetylation and H3K9me3, which marks heterochromatic regions in the genome (87) decrease upon SIRT5 loss. Likewise, oxidative stress increases in SIRT5-depleted melanoma cells, consistent with impaired regeneration of reduced glutathione, a major antioxidant species and an output of 1CM.

Many open questions remain as to the mechanisms by which SIRT5 promotes proper gene expression and viability in melanoma. The accumulation of acetyl-CoA in SIRT5-depleted melanoma cells suggests that SIRT5 may promote the activity of a histone acetyltransferase to promote histone acetylation, a possibility that we are currently investigating. Alternatively, SIRT5 could promote generation of a localized acetyl-CoA pool necessary to drive histone acetylation (*cf.* the nuclear pool (74)), without influencing global acetyl-CoA levels. A large number of studies implicate alterations in levels of specific metabolites in driving chromatin modifications (88). Increased lactate, for example inhibits histone deacetylases, thereby increasing histone acetylation (89). Although we observe only modest and, in some cases, inconsistent changes in cellular metabolite levels upon SIRT5 KD, it is possible that alterations in levels of specific metabolites, or a combination of these metabolite abnormalities, may in part be responsible for the loss of histone modifications we observe. In addition, we identified MTHFD1L as a novel SIRT5 interactor and candidate target that may play a role in SIRT5-mediated regulation of 1CM. Unfortunately, we have been unsuccessful at rescuing the cellular lethality associated with SIRT5 depletion using relevant small molecule metabolites or drugs (acetate, acetyl-CoA, SAM, serine, glycine, histone deacetylase and demethylase inhibitors, antioxidants, nucleotides, and amino acids [data not shown]). We suspect that this reflects pleiotropic functions and targets of SIRT5 in melanoma cells, impairment of which cannot be rescued by intervention in any individual pathway. SIRT5 targets involved in other pathways -- *e.g.* ROS suppression, cell death (32, 90), and others -- could well contribute to the requirement of melanoma cells for SIRT5. Likewise, we identified perturbations in innate immune pathways in SIRT5-depleted melanoma cells, which could also contribute to the requirement of melanoma cells for this protein. This is consistent with the hundreds of cellular targets of SIRT5, involved in diverse cellular pathways, identified in proteomics studies (17). Moreover, it is consistent with the observation that SIRT5 plays pro-survival roles across multiple different cancer types, via distinct proposed mechanisms. As the dominant cellular desuccinylase/demalonylase/deglutarylase, it is possible that SIRT5 is recruited to play distinct roles in supporting tumorigenesis, modulating activities of different suites of targets and pathways, in a cancer type-specific manner.

Overall, our data reveal a major, hitherto unknown requirement for SIRT5 in melanoma cell survival, through suppression of apoptosis via regulation chromatin modifications and expression of critical pro-survival genes, including MITF and c-MYC (Figure 8G). These results, along with those already in the literature (7), suggest that SIRT5 may play potent oncogenic roles across many diverse tumor types, seemingly engaging a variety of different cellular mechanisms to do so in a cancer- and context-specific manner. Since the phenotypes of *Sirt5* null mice are quite mild, we propose that SIRT5 may represent an attractive new therapeutic target, in melanoma and specific other cancer types. In this regard, published studies (17, 91–94), including recent work focused on breast cancer (37) demonstrate that SIRT5 is in principle druggable with small molecules. SIRT5 dependency may be particularly translationally significant in uveal melanoma, where currently no effective therapeutic options exist for patients with metastatic disease.

## Materials and Methods

### Analysis of SIRT5 gene amplification, mRNA and protein expression in melanoma

Percentages of genetic alterations, as indicated, in *SIRT1-SIRT5*, *BRAF*, *NRAS*, *PTEN*, *MITF* and *NF1* in human melanoma cases were calculated from TCGA data where copy number and mutation data were available (n=287, Provisional, analyzed on cBioPortal). Kaplan–Meier analysis of overall or disease-free survival in melanoma patients with or without copy number gain or amplification of *SIRT5* were similarly analyzed. Sirtuin copy number analysis of melanoma cell lines was performed by high density SNP array of 8 primary, 72 stage III, 51 stage IV and 8 stage III/IV (metastatic disease) melanoma cell lines, as previously described (48, 49). Data were analyzed using Nexus Copy Number (BioDiscovery) for copy gain, copy loss and loss of heterozygosity of genes and chromosomal regions. Data from the TCGA was used via cBioPortal to further investigate *SIRT5* in melanoma (n=478; Cancer Genome Atlas Network), including copy number (by GISTIC 2.0) and expression data (by RNAseq) and correlation with clinical attributes, including the Clark level at diagnosis of the melanoma.

### Immunohistochemical analysis of SIRT5 in human nevus and melanoma

The study was undertaken with Human Ethics Review Committee approval and patient’s informed consent. The Melanoma Institute Australia Medical Research Database and archival files of the Department of Tissue Pathology and Diagnostic Oncology, Royal Prince Alfred Hospital, were utilized to identify melanocytic lesions (human ethics committee approval X11-0289, HREC/11/RPAH/444). Melanocytic lesions were selected based on the pathological diagnosis of the melanocytic lesion as a compound nevus, dysplastic nevus, thin primary melanoma (Breslow thickness <1mm), thick primary melanoma (Breslow thickness >1mm) or metastatic melanoma in regional lymph node or distant metastatic site. Immunohistochemical SIRT5 staining intensity in melanocytes was scored by two pathologists blinded as to the diagnosis associated with each tissue core. Scores were then averaged.

### Cell culture

Cutaneous melanoma cell lines with mutations in either *NRAS* or *BRAF*: A375, A2058, SK-MEL-2, SK-MEL-28, VMM15, and VMM917 were purchased from ATCC (Table S2); SK-MEL-19 was generously provided by Monique Verhaegen (UM); C8161 was provided by Dr. Zaneta Nikolovska-Coleska (UM); SK-MEL-239 and vemurafenib-resistant (SK-MEL-239**VR)** lines were generously provided by Dr. Emily Bernstein (ISMMS). Uveal melanoma cell lines MP-38, MP-41 and MP-46 were purchased from ATCC; cell line 92-1 was purchased from Sigma-Aldrich (Table S2). Ovarian cancer cell lines A2780, SK-OV-3, OVCAR4 and OVCAR10 were generously provided by Kathleen Cho (UM).

Unless otherwise noted, A375, A2058 and SK-MEL-19 cell lines were cultured in DMEM (Gibco) containing 4.5g/L glucose, 110mg/L sodium pyruvate, 4mM L-glutamine, 100units/mL penicillin, 100µg/mL streptomycin and 10% heat-inactivated FBS. SK-MEL-2 and SK-MEL-28 cell lines were cultured in MEM (Gibco) containing 1.0g/L glucose, 110mg/L sodium pyruvate, 2mM L-glutamine, 0.1mM non-essential amino acids (Gibco), 100units/mL penicillin, 100µg/mL streptomycin solution and 10% heat-inactivated FBS. VMM15, VMM917, SK-MEL-239 and SK-MEL-239**VR** cell lines were cultured in RPMI (Gibco) containing 4.5g/L glucose, 10mM HEPES, 110mg/L sodium pyruvate, 4mM L-glutamine, 100units/mL penicillin, 100µg/mL streptomycin and 10% heat-inactivated FBS. SK-MEL-239**VR** derivative cells were cultured in 2µM vemurafenib (Cayman Chemical). The C8161 cell line was grown in DMEM/F12 (1:1) (Gibco) containing 1.2mM L-glutamine, 0.1mM non-essential amino acids, 100units/mL penicillin, 100µg/mL streptomycin and 5% heat-inactivated FBS. MP-38, MP-41 and MP-46 cells were cultured in RPMI (Gibco) containing 2 mM L-glutamine, 10 mM HEPES, 110mg/L sodium pyruvate, 4.5g/L glucose, 100units/mL penicillin, 100µg/mL streptomycin and 25% heat-inactivated FBS; 92-1 cells were cultured in RPMI (Gibco) containing 2 mM L-glutamine, 10 mM HEPES, 110mg/L sodium pyruvate, 4.5g/L glucose, 100units/mL penicillin, 100µg/mL streptomycin and 10% heat-inactivated FBS. S-adenosylmethionine (SAM) resuspended in 0.005M sulfuric acid and 10% ethanol was purchased from New England Biolabs. All cell lines were routinely confirmed to be free of mycoplasma contamination by PCR assay and were grown in a humidified chamber at 37°C containing 5% CO_2_. All cutaneous melanoma cell lines were authenticated via STR profiling at the University of Michigan DNA Sequencing Core (data available upon request). Table S2 lists the genetic alteration of each cell line, and the sex and age of the patient at the time of cell line derivation, if known. Ovarian cancer cell lines A2780, OVCAR10, OVCAR4 were cultured in RPMI (Gibco) containing 2 mM L-glutamine, 10 mM HEPES, 110mg/L sodium pyruvate, 4.5g/L glucose, 100units/mL penicillin, 100µg/mL streptomycin and 10% heat-inactivated FBS. SK-OV-3 cells were cultured in McCoy’s (Gibco) medium supplemented with 100units/mL penicillin, 100µg/mL streptomycin and 10% heat-inactivated FBS.

### Lentiviral transduction

Lentiviral plasmids in the pLKO.1 backbone, encoding puromycin *N*-acetyl-transferase, containing shRNAs targeting human *SIRT5* or murine *Sirt5* (Table S6) were used to generate high-titer lentiviral particles at the Vector Core (UM). A non-silencing control shRNA against Gaussia luciferase was used as the non-targeting control (NT). KD3 and KD4 shRNA lentiviral plasmids targeting human *SIRT5* were designed based on previously published CRISPR screens (95, 96). All other shRNA plasmids were purchased through the Vector Core (UM). Lentiviral transduction was carried out in the presence of 8 µg/ml polybrene in complete growth medium for 24 hrs., after which medium was replaced with fresh complete growth medium. Puromycin was added to a final concentration of 1 µg/ml 48 hrs. post-transduction to select for transductants. Successful SIRT5 KD was routinely confirmed by western blotting 48-96 hrs. post-transduction.

### Proliferation assays

Forty-eight hrs. after lentiviral transduction, 5×10^3^ cells were plated into 96-well plates in the presence of 1µg/ml puromycin or in 1µg/ml puromycin with a cocktail of 0.1mM non-essential amino acids (Gibco) and 5mM alpha-ketoglutarate (Sigma), where indicated. Twenty-four hrs. after plating, relative cell mass was assessed using WST-1 Cell Proliferation Reagent (Clontech) per manufacturer’s instruction. After addition of WST-1, plates were incubated at 37°C for 2 hrs. before reading the optical density at 450nm. OD450nm was assessed every 24 hrs. as indicated and plotted.

Ovarian cancer cell lines, A2780, SK-OV-3, OVCAR10 and OVCAR4, were transduced with lentiviral particles carrying either a non-silencing control shRNA against Gaussia luciferase, KD1 or KD2 shRNA targeting human *SIRT5*. Lentiviral transduction was carried out in the presence of 8 µg/ml polybrene in complete growth medium for 24 hrs., after which medium was replaced with fresh complete growth medium. Puromycin was added to a final concentration of 2 µg/ml 48 hrs. post-transduction to select for transductants. Successful SIRT5 KD was confirmed by western blotting. Following puromycin selection and confirmation of SIRT5 KD, 2.5×10^4^ cells were plated in complete growth media in triplicates into 12-well plates. Cell number in each well were assessed every 24 hrs. after plating by manual counting using a hemocytometer.

### Colony formation assays

Two million A2058 or SK-MEL-2 melanoma cell lines were lentivirally transduced in a 10-cm dish with a non-silencing shRNA or one of two shRNAs targeting human *SIRT5*. Forty-eight hrs. post-transduction 1×10^5^ cells were plated into each of four wells of a six-well dish as previously described. Twelve days after transduction, puromycin-selected cells were stained with 0.25% (w/v) crystal violet (Sigma) in 20% ethanol for 30 minutes according to standard protocols. Colony formation was quantified by solubilizing crystal violet in a 30% methanol, 10% acetic acid solution and absorbance measured at OD590nm.

### CRISPR/Cas9 targeting of SIRT5

CRISPR plasmids were generated as described (97). Briefly, guide sequences (Table S6) targeting the *SIRT5* locus were inserted into pSpCas9(BB)-2A-Puro (PX459) backbone (Addgene catalog #62988). Guide sequences were designed based on previous CRISPR screens (95, 96, 98). Sanger sequencing confirmed successful cloning of guide sequences. A2058 cells were seeded at 50,000 cells/well two days prior to transfecting 500ng of each indicated plasmid using polyethylenimine (PEI) in DMEM without serum or antibiotics. Transfected cells were selected using puromycin (1 µg/ml) for 7 days, beginning 24 hrs. post-transfection. Fourteen days after transfection, cells were fixed and stained with 0.25% crystal violet in 20% ethanol for 30 mins at room temperature. Colony formation was quantified by solubilizing crystal violet in a 30% methanol, 10% acetic acid solution and absorbance measured at OD590nm. Confirmation of SIRT5 KD via immunoblot was performed 33 days post-transfection.

### Immunoblotting

Whole-cell protein extracts were prepared in protein sample buffer (62.5mM Tris pH 6.8, 2% SDS, 10% glycerol). Lysates were sonicated for 30 seconds using a Branson Sonifier set to output “2.” Lysates were then clarified by centrifugation at 15,000rcf for 30 minutes at 4°C. Protein concentrations were determined using DC Protein Assay (Bio-Rad). Equivalent amounts (10-50 µg) of total protein, supplemented with 710mM β-mercaptoethanol and 0.01% (w/v) bromophenol blue, then boiled for 5 minutes, were fractionated by SDS-PAGE on a 12% or 15% polyacrylamide gel, electrophoretically transferred to PVDF, and probed with antibodies diluted in 5% nonfat milk in 1XTBS-0.1% Tween-20. Membranes were imaged on an ImageQuant LAS 4000 Scanner (GE Healthcare) after application of Immobilon Western HRP Substrate (Millipore). See Table S7 for antibodies used in this study.

### Immunofluorescence

A2058 cells were plated at a density of 0.5×10^6^ per well in six-well plates containing glass coverslips (Werner #1.5 thickness, 12mm diameter). Twenty-four hrs. later, cells were transduced with either a lentivirus expressing a non-silencing shRNA (control) or one of two shRNAs targeting *SIRT5* (KD1 or KD2). Ninety-six hrs. after transduction, puromycin-selected cells were incubated with 100nM Mitotracker (BD Biosciences) for 30 minutes at 37°C, washed with PBS, and fixed in 3.7% formaldehyde. After permeabilization in 0.3% Triton X-100 for 10 minutes at room temperature, cells were blocked with 5% normal goat serum in 0.2% Triton X-100 in PBS for one hour. Cells were incubated in SIRT5 (Sigma) primary antibody diluted to 1 µg/ml in 5% bovine serum albumin in 0.2% Triton X-100 in PBS overnight at 4°C. Cells were then washed 3 times in 1X PBS and incubated in Alexa 488 anti-rabbit (Invitrogen) diluted 1:1000 in 5% bovine serum albumin in 0.2% Triton X-100 in PBS for 1 hour at room temperature. Cells were washed 2 times in 1X PBS, and 1X with PBS supplemented with 0.01 µg/ml DAPI. Cells were mounted with prolong gold antifade reagent (ThermoFisher) and imaged on an Olympus FV 500 Confocal microscope.

### Cellular Fractionation

Lentivirally-transduced A2058 cells were washed twice with 1X PBS and harvested by scraping 96hrs. post-transduction. Subcellular fractionation was carried out using the CelLytic NuCLEAR Extraction Kit (Sigma-Aldrich), according to the manufacturer’s protocol. The cytoplasmic fraction was concentrated using Amicon Ultra 0.5 mL Centrifugal Filters (Sigma-Aldrich), according to manufacturer’s instructions. Each fraction and whole-cell pellets were resuspended in protein sample buffer, processed as previously (see above), and analyzed by immunoblot.

### Annexin V flow cytometry

A2058 and SK-MEL-2 cells were plated at a density of 0.5×10^6^ per well of two six-well plates/cell line. Four wells of each cell line were transduced with either a lentivirus expressing a non-silencing shRNA (control) or one of two shRNAs targeting *SIRT5* (KD1 or KD2). Ninety-six hrs. after transduction, puromycin-unselected cells were harvested and stained for 30 minutes at room temperature with FITC Annexin V (BD Biosciences) and propidium iodide (Sigma), according to the BD Biosciences staining protocol. Cells were analyzed by flow cytometry using a BD FACSCalibur and results were plotted using FlowJo 10.2 analysis software.

### Xenograft assays

A2058 puromycin-unselected cells were harvested 72 hrs. post-transduction with pLKO control, pLKO *SIRT5* KD1, or pLKO *SIRT5* KD2. Subcutaneous tumor growth was initiated by injection of 1×10^6^ cells of each cell line, resuspended in 1:1 DMEM:Matrigel Matrix (Corning), into the contralateral flanks of 11-13 week old NOD.Cg-*Prkdc^scid^Hr^hr^*/NCrHsd female mice (Envigo). Each experimental group contained 5 mice. Tumor size was measured in millimeters (mm) using Vernier calipers at the timepoints indicated. Tumor volume was calculated according to the formula:

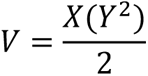

where *V* is tumor volume in mm^3^, *X* is the longest length of the tumor, and *Y* is the shortest length of the tumor, perpendicular to *X* (99, 100). All mice were euthanized when a tumor ulcerated or reached 2000mm^3^. All mice were housed at the Biomedical Science Research Building (UM). All vertebrate animal experiments were approved by and performed in accordance with the regulations of the University Committee on Use and Care of Animals.

### Tumor induction in Braf^CA^;Pten^fl/fl^;Tyr::CreER;Sirt5 mice

*Sirt5* KO mice (15) were crossed with *Braf^CA^;Pten^fl/fl^;Tyr::CreER* mice (55) to generate groups of littermate *Braf^CA^;Pten^fl/fl^;Tyr::CreER;Sirt5* KO and WT controls on a mixed BL/6/129SvJ genetic background. For melanoma induction, flanks of adult mice (4-9 weeks of age) were treated with depilatory cream with a cotton applicator. After 3-5 minutes, the area was rinsed with distilled water, and the treatment spot marked. Topical administration of 4-hydroxytamoxifen (4-HT; 25mg/ml in DMSO, Cayman Chemical) was repeated on each of three consecutive days, by applying one microliter of 4-HT to the spot. Mice were monitored for tumor appearance, which occurred typically within 4-8 weeks, and euthanized when a tumor ulcerated or reached 2000mm^3^. All mice were housed at the Biomedical Science Research Building (UM). Experiments were approved by and performed in accordance with the regulations of the University Committee on Use and Care of Animals.

### Transcriptomic analysis of SIRT5 depletion

A2058, A375 and SK-MEL-2 cells were plated at a density of 0.5×10^6^/well of two six-well plates per cell line. Four wells of each cell line were transduced with either a lentivirus expressing a non-silencing shRNA (control) or one of two shRNAs targeting *SIRT5* (KD1 or KD2). Total RNA was extracted in TRIzol (Invitrogen) 96 hrs. post-transduction from three wells and treated with RNAse-free DNAse I (Roche) for 1 hour at 37°C, according to manufacturer’s instructions. RNA samples (n=3 biological replicates for each of 3 cell lines for each condition) were submitted to the University of Michigan Sequencing Core for sample processing and Illumina HiSeq-4000 50nt paired-end sequencing. The remaining well was harvested for protein to confirm SIRT5 KD by immunoblot. Illumina libraries were prepared using random primers according to manufacturer’s instructions.

Paired-end reads were assessed for various quality metrics before and after trimming via FastQC v.0.11.9 (101) and MultiQC v.1.9 (102). All quality control modules investigated suggested high sample quality and removal of adapter content. Reads were trimmed by Trimmomatic v.0.39 (103) in paired-end mode, yielding filtered R1 and R2 FASTQ files containing only mate-paired reads. Trimmed FASTQ files were aligned by STAR v.2.7.3a (104) in paired-end mode against Gencode (GRCh38 Release 34 primary assembly). For each unique sample, all corresponding FASTQ files (i.e. R1/R2, split lanes) were aligned together in a single STAR call via –readFilesIn to yield single sorted BAM files for each sample. STAR-output BAM files were subsequently counted by the SubRead v.2.0.0 (105) featureCounts command to yield counts. Count matrices produced by featureCounts were then supplied to the Bioconductor package DESeq2 v.1.28.1 (106) for differential gene expression analysis. Contrasts controlling for cell line and pooling unique knockdowns together were utilized to discern the generalized impacts of SIRT5 knockdown. DESeq2 results were first independently filtered, and then the unique, non-censored genes across all contrasts of interest were combined to generate a standardized list of genes that were re-evaluated for differential expression without independent filtering to ensure gene-level statistics were generated for every contrast and not missing. All log2(Fold Change) values were subsequently shrunk, utilizing the apeglm (v.1.10.0) package’s “Approximate Posterior Estimation for GLM” (apeglm) algorithm (107), providing Bayesian shrinkage estimators for the effect size. Genes with calculated p-values, after adjusting for multiple comparisons by the Benjamini–Hochberg method (i.e. q-values), at p<0.01 (5% FDR) and a |log2(Fold Change)|>1 were then considered significant. FPKM values were estimated using DESeq2 FPKM function in robust size factor normalization mode.

### Differently expressed gene correlation with SIRT5 expression in human melanoma

TCGA RNA data (EBPlusPlusAdjustPANCAN_IlluminaHiSeq_RNASeqV2.geneExp.tsv) were downloaded from the Pan-Cancer Atlas site (PanCanAtlas; https://gdc.cancer.gov/about-data/publications/pancanatlas; accessed November, 2020, (108)). These data were filtered to encompass 474 sequenced human skin cutaneous melanoma samples with RNA abundance data (i.e. counts). The counts of differently expressed genes (DEGs) in response to aggregate SIRT5 KD (q<0.05) that uniquely mapped to Entrez Ids were correlated with SIRT5 counts in these clinical sample via Spearman’s rank correlation. Genes of interest were identified by overlapping genes evaluated for correlation with known oncogenes from the ONGene database ((109); http://ongene.bioinfo-minzhao.org/) and melanoma subtype signatures defined previously (110). Prior to intersection, the melanoma subtype signatures (gene symbols) were updated using the HGNC Multi-symbol checker (https://www.genenames.org/tools/multi-symbol-checker/) to maximize the number of genes overlapping those in the more recent genome annotation we utilized.

### Code Availability

Supporting analysis code, files, and analysis documentation is hosted at https://github.com/monovich/giblin-sirt5-melanoma.

### SIRT5 RNA-seq gene ontology

Differentially expressed genes were classified by a padj-value<0.05. To identify shared or unique protein-coding genes between A2038, A375, and SK-MEL-2 datasets, differentially expressed protein-coding genes from each cell line were first designated as up- or down-regulated upon SIRT5 KD. Each up- or down-regulated protein-coding list was then cross-referenced between the three cell lines to generate the values indicated in the Venn diagram. Gene set enrichment analysis (GSEA (111)) was performed using the Preranked function in the GSEA 4.1.0 software against the MSigDB 7.2 release (112) on the unfiltered RNAseq dataset of pooled SIRT5 KD against pooled control samples (GSEA references here). The list of ranked genes used as input for GSEA analysis was generated by multiplying the sign of fold-change (+ or -) with -log(padj). Additional gene ontology was performed using Ingenuity Pathway Analysis (IPA: QIAGEN Inc.) on transcripts with a padj-value<0.05 from the RNAseq dataset of pooled SIRT5 KD against pooled control samples. All pathways with a p-value<0.05 (-log(p-value)>1.3) were deemed significant. As defined by IPA, significant upstream regulators were defined as significant based on a p-value<0.05 and an absolute activation z-score greater than 2.

### CUT&RUN

CUT&RUN for H3K9ac in A2058 cells was carried out using the CUT&RUN Assay Kit (Cell Signaling Technology), according to the manufacturer’s instructions. Briefly, 2×10^5^ A2058 cells were harvested 96hrs. after transduction with either a lentivirus expressing a non-silencing shRNA (control) or one of two shRNAs targeting *SIRT5* (KD1 or KD2) and subjected to chromatin immunoprecipitation using 5µl of either anti-H3K9ac or, as a negative control, rabbit IgG isotype antibody. Samples were incubated overnight with shaking at 4°C. DNA was then extracted via ethanol precipitation overnight at −20°C, and then resuspended in 50µl of water. The purified DNA (2µl per reaction) was quantified by real-time PCR using SimpleChIP human *MITF* (Cell Signaling Technology) or human *c-MYC* primers (forward: 5’-GGACCCGCTTCTCTGAAAGG-3’ and reverse: 5’-GCAAGTGGACTTCGGTGCTTACC-3’, as previously described (113)), and SYBR Select Master Mix (Applied Biosystems). Ct values were normalized to control H3K9ac values. ChIP-qRTPCR was done three independent times with n=3 technical replicates each time.

### Identifying differentially active reactions using genome-scale metabolic modeling

Gene expression data from the melanoma cell lines A2058, A375, and SK-MEL-2 was used as input to identify differentially active reactions in the human genome-scale metabolic model using a modeling approach detailed in Shen *et al.* (114, 115). This approach identifies a metabolic flux state that best fits the transcriptomics profile in each condition. Normalized expression data for each cell line was compared between the two groups (C vs. KD) to attain a list of significantly up- and down-regulated genes using a significance threshold of p<0.05 and a z-score threshold above 2 or below −2, respectively. These genes were then overlaid onto the metabolic model based on gene-protein-reaction annotations in the model. Finally, reaction flux data was generated using a linear optimization version of the iMAT algorithm (71, 115) with the following inputs: the RECON1 model, the list of up- and down-regulated genes, and the recommended values for the optional parameters (rho = 1E-1, kappa = 1E-1, epsilon = 1, mode = 0). This resulted in flux predictions for all three cell lines. Differentially active reactions were identified by transforming the fluxes into z-scores and estimating the significance of the difference between z-scores before and after KD across all cell lines using a t-test. The flux data for all the reactions is available in Table S4. Flux data was visualized through heatmaps generated using R software, where reactions were clustered by rank correlation. Only reactions with significant flux difference between control and KD groups (p<0.01) are shown. Transport reactions were excluded from the analysis.

### Acetyl-CoA Quantification

Acetyl-CoA were quantified by stable isotope dilution liquid chromatography-high resolution mass spectrometry, as previously described (75, 76). Cell pellets were spiked with an internal standard prepared as described (75), and sonicated for 12 cycles of 0.5 sec. pulses in 10% (w/v) trichloroacetic acid (Sigma Aldrich) in water. Protein was pelleted by centrifugation at 17,000rcf for 10 min. at 4°C. The cleared supernatant was purified by solid-phase extraction using Oasis HLB 1cc (30 mg) SPE columns (Waters). Columns were washed with 1ml methanol, equilibrated with 1ml water, loaded with sample, desalted with 1ml water, and eluted with 1ml methanol containing 25 mM ammonium acetate. The purified extracts were evaporated to dryness under nitrogen and resuspended in 55μl 5% (w/v) 5-sulfosalicylic acid (SSA) in optima HPLC grade water. Acetyl-CoA was measured by liquid chromatography-high resolution mass spectrometry. Briefly, 5μl of sample in 5% SSA were analyzed by injection into an Ultimate 3000 HPLC coupled to a Q Exactive Plus (Thermo Scientific) mass spectrometer in positive ESI mode using the settings described previously (76). Calibration curves were prepared using analytical standards from Sigma Aldrich and processed identically as the samples. Data were integrated using Tracefinder v4.1 (Thermo Scientific) software, and additional statistical analysis conducted by Prism v7.05 (GraphPad). Acetyl-CoA values were normalized to cell number and reported as pmol/10^5^ cells.

### Targeted metabolomics analysis of SIRT5 depletion

Melanoma cell lines infected with a control (pLKO.1 empty vector), SIRT5 KD1 or KD2 virus were cultured for 72 hrs. in complete growth medium on 10cm dishes in biological sextuplicate. Three dishes were reserved for metabolite collection, and three dishes were harvested in protein sample buffer (62.5mM Tris pH 6.8, 2% SDS, 10% glycerol) for protein quantification. A complete medium change was performed two hrs. prior to metabolite collection. Cells were washed twice in 2ml of PBS; one wash at room temperature and one at 4°C. Taking care to aspirate all remaining liquid, plates were placed onto dry ice to cool before incubating in 4ml of 80% methanol for 10 mins. Plates were then scrape-harvested, and lysates transferred to 15 ml conical tubes. Lysates were centrifuged at 200 x g for 10 mins. For all experiments, the quantity of the metabolite fraction analyzed was adjusted to the corresponding average protein concentration. Liquid chromatography coupled-tandem mass spectrometry (LC-MS/MS) was employed for the detection of relative metabolite levels (116). Samples were analyzed on a 6490 Triple Quadrupole (QqQ) LC-MS against a targeted panel of 225 metabolites run in positive and negative modes. Agilent MassHunter Optimizer and Workstation Software LC-MS Data Acquisition for 6400 Series Triple Quadrupole B.08.00 was used for standard optimization and data acquisition. Agilent MassHunter Workstation Software Quantitative Analysis Version B.0700 for QqQ was used for initial raw data extraction and analysis. Each MRM transition and its retention time of left delta and right delta was 1 min. Additional parameters include mass extraction window of 0.05 Da right and left from the extract m/z, Agile2 integrator algorithm, peak filter of 100 counts, noise algorithm RMS, noise SD multiplier of 5 min, S/N 3, Accuracy Max 20% max %Dev, and Quadratic/Cubic Savitzky-Golay smoothing algorithm with smoothing function width of 14 and Gaussian width of 5. Peak area values under 10,000 were discarded as noise. Remaining raw values for each metabolite were median-centered across all conditions for each cell line. The average and standard deviation were taken for all replicates, and Student’s t-tests were conducted comparing each KD to the control. Data for each metabolite was represented as log(2) fold change with relative standard deviation. Pathway enrichment studies of both up- and down-regulated signaling were completed using Metaboanalyst and represented as log10(p-value) (117).

### CM-H2DCFDA staining for ROS measurement

A2058 cells were plated at a density of 1×10^6^ per well of two six-well plates. Three wells were transduced each with either a lentivirus expressing a non-silencing shRNA (control) or one of two shRNAs targeting *SIRT5* (KD1 or KD2). Ninety-six hrs. after transduction, CM-H2DCFDA (ThermoFisher) was added to a final concentration of 5μM and incubated for 30 minutes in a humidified chamber at 37°C. Cells were analyzed by flow cytometry using a BD FACSCalibur and results were plotted using FlowJo 10.2 analysis software.

### Co-immunoprecipitation of SIRT5 with MTHFD1L

A2058 whole-cell protein extracts were prepared in protein lysis buffer (50mM Tris pH 7.4, 150mM NaCl, 1% Triton X-100, 0.5% NP40, 10% Glycerol) supplemented with 1µM TSA, 10mM nicotinamide, PhosStop cocktail (Roche), and protease inhibitor cocktail (Roche). Lysates were sonicated for 30 seconds using a Branson Sonifier set to output “2.” Lysates were then clarified by centrifugation at 15,000rcf for 30 minutes at 4°C. Protein concentrations were determined using DC Protein Assay (Bio-Rad). Equivalent amounts (1mg) of total protein were mixed with the indicated amount (or 5μg) of biotinylated anti-SIRT5 antibody or normal rabbit IgG, and 20µl of protein A magnetic beads (Pierce), equilibrated in lysis buffer. After overnight rotation at 4°C, samples were washed 4 times in 1ml of lysis buffer. Following the final wash, samples were boiled in 62.5mM Tris pH 6.8, 2% SDS, and 10% glycerol, supplemented with 710mM β-mercaptoethanol and 0.01% (w/v) bromophenol blue for 5 mins, and fractionated by SDS-PAGE, followed by electrophoretic transfer and immunoblotting with the indicated antibodies.

### Measurement of Oxygen consumption, Extracellular acidification, and Real-Time ATP production rate

Oxygen consumption rate (OCR), Extracellular acidification rate (ECAR), and Real-Time ATP production rate were measured using the XFe96 Extracellular Flux Analyzer (Seahorse Bioscience, Agilent Technologies, Santa Clara, CA). To measure glucose-dependent mitochondrial respiration, mitochondrial stress tests were performed to measure OCR per manufacturer’s instructions. Briefly, 72 hrs. post-transduction, 4×10^4^ A375 or A2058 cells were plated in DMEM complete media supplemented with 10% heat-inactivated FBS into each well of a 96-well Seahorse microplate. Cells were then incubated in 5% CO_2_ at 37°C for 24 hrs. Following incubation, cells were washed twice, incubated in non-CO_2_ incubator at 37°C, and analyzed in XF assay media (non-buffered DMEM containing 25mM glucose, and 1mM sodium pyruvate, pH 7.4) at 37°C, under basal conditions and in response to 2μM oligomycin (Sigma), 1μM fluoro-carbonyl cyanide phenylhydrazone (FCCP) (Sigma) and 0.5μM rotenone (Sigma)/0.5μM antimycin A (Sigma). Data were analyzed by the Seahorse XF Cell Mito Stress Test Report Generator. OCR (pmol O_2_/min) values were normalized to the protein content.

To measure glutamine-dependent mitochondrial respiration, mitochondrial stress test was performed to measure OCR as per manufacturer’s instructions. Briefly, 72 hrs. post-transduction, 4×10^4^ A375, A2058 or SK-MEL-2 cells were plated in DMEM complete media supplemented with 10% heat-inactivated FBS into each well of a 96-well Seahorse microplate. Cells were then incubated in 5% CO_2_ at 37°C for 24 hrs. Following incubation, cells were washed twice, incubated (in non-CO_2_ incubator at 37°C), and analyzed in XF assay media (non-buffered DMEM containing 4 mM L-glutamine, and 1 mM sodium pyruvate, pH 7.4) at 37°C, under basal conditions and in response to 2μM oligomycin (Sigma), 1μM fluoro-carbonyl cyanide phenylhydrazone (FCCP) (Sigma) and 0.5μM rotenone (Sigma)/0.5μM antimycin A (Sigma). Data were analyzed by the Seahorse XF Cell Mito Stress Test Report Generator. OCR (pmol O_2_/min) values were normalized to protein content.

ECAR values were measured by performing glycolysis stress tests according to manufacturer’s instructions. Briefly, 72 hrs. post-transduction, 4×10^4^ A375 and A2058 cells were plated in DMEM complete media supplemented with 10% heat-inactivated FBS into each well of a 96-well Seahorse microplate. Cells were then incubated in 5% CO_2_ at 37°C for 24 hrs. Following incubation, cells were washed twice, incubated in a non-CO_2_ incubator at 37°C, and analyzed in XF assay media (non-buffered DMEM containing 2mM L-glutamine, pH 7.4) at 37°C, under basal conditions and in response to 10mM glucose (Sigma), 2μM oligomycin (Sigma), and 50mM 2-deoxy-D-glucose (Sigma). Data were analyzed by the Seahorse XF Cell Glycolysis Stress Test Report Generator. ECAR (mpH/min) values were normalized to protein content.

The Seahorse XF Real-Time ATP Rate Assay Kit (Agilent) was used to simultaneously measure the basal ATP production rates from mitochondrial respiration and glycolysis. The assay was performed per manufacturer’s instructions. Briefly, 72 hrs. post-transduction, 4×10^4^ A375 and A2058 cells were plated in DMEM complete media supplemented with 10% heat-inactivated FBS into each well of a 96-well Seahorse microplate. Cells were then incubated in 5% CO_2_ at 37°C for 24 hrs. Following incubation, cells were washed twice, incubated in a non-CO_2_ incubator at 37°C, and analyzed in XF assay media (non-buffered DMEM containing 10mM glucose, 2mM L-glutamine, and 1mM sodium pyruvate, pH 7.4) at 37°C, under basal conditions and in response to 1.5μM oligomycin, and 0.5μM rotenone/0.5μM antimycin A. Data were analyzed by the Seahorse XF Real-Time ATP Rate Assay Report Generator. ATP production rates were normalized to protein content.

### JC-1 Mitochondrial membrane potential assay

At 96 hrs. post-transduction approximately 2×10^5^ A2058 cells in 100µl warm PBS were transferred into tube and stained with 10 µg/ml of JC-1 dye (AnaSpec). An additional 100µl from control cells were treated with 200 µM FCCP, a mitochondrial uncoupler, which depolarizes mitochondrial membrane potential, and used as a positive control. After 30 minutes of staining, cells were centrifuged to remove excess JC-1, washed twice with warm PBS and resuspended in 100µl of warm PBS. Cells were transferred into black 96-well plates and absorbance measured at 535nm (excitation)/595nm (emission) (aggregate, red) and 485nm (excitation)/435nm (emission) (monomer, green) using a H1 Synergy plate reader. The data are presented as a ratio of red to green fluorescence.

### [^13^C_6_] Glucose, [^13^C_5_] Glutamine and [^15^N_2_] Glutamine Labelling for Metabolic Flux

Glucose and glutamine labelling were carried out as described previously (58). Briefly, 72 hrs. after transduction with pLKO control, pLKO *SIRT5* KD1, or pLKO *SIRT5* KD2, cells were trypsinized and plated in triplicate in 6-well dishes at a density of 1×10^6^ cells per well. Culture medium was replaced with [^13^C_6_] glucose, [^13^C_5_] glutamine or [^15^N_2_] glutamine labelling medium for 6 hours. Glucose labeling medium consisted of: MEM (1 g/L glucose; Gibco) supplemented with 1 g/L [U-^13^C_6_] glucose (Sigma), 10% v/v fetal bovine serum, 2 mM L-glutamine, 1% v/v pen/strep solution, and 1% MEM non-essential amino acids. Glutamine labeling medium was prepared as the glucose labeling medium, except an additional 1 g/L of unlabeled glucose, 1 mM unlabeled L-glutamine, and 1 mM [U-^13^C_5_] L-glutamine (Sigma) or 1 mM [^15^N_2_] L-glutamine (Sigma) was added. After 6 hours, cells were washed with cold PBS, then 0.45 ml of 50% methanol:50% water containing the internal standard, 20 µM L-norvaline (Sigma) was added to each well. Plates were frozen on dry ice for 30 minutes and thawed on ice for 10 minutes. The cell suspension was transferred to a microfuge tube followed by the addition of 0.225 ml chloroform. Samples were vortexed, then centrifuged at 20,000rcf for 5 minutes at 4°C. The top layer was transferred to a fresh tube and dried in a Speedvac. Samples were then shipped to the Sanford Burnham Prebys Medical Discovery Institute (La Jolla, CA) for GC/MS-based metabolic profiling. Derivatization of metabolites, GC/MS settings, and data analysis for stable isotope labeling and metabolite quantification were as described (61). Metabolite quantities were determined using mass ion peak areas corresponding to unlabeled metabolites, and then corrected for the fraction of a metabolite that was ^13^C or ^15^N labeled to yield the total (labeled plus unlabeled) cellular quantity for that metabolite. This amount was normalized to total cellular protein.

### Statistical Analysis

All statistical analyses were performed using Prism graphing software (Graphpad). Unless otherwise noted, p≤0.05 produced from an unpaired Student’s t-test was considered significant.

### Study approval

All mice were housed at the Biomedical Science Research Building (UM). All vertebrate animal experiments were approved by and performed in accordance with the regulations of the University Committee on Use and Care of Animals.

## Author Contributions

WG, LBR, AHG, SK, ACM, AMM, MES, MA, ASAM, CHC, NK, KAM, HJL, LZ, PS, ST, ELV, SI, MW, JSW, HPS, RAS, ALP, AA, RAS, MSS, DAS, DRF, MWB, SC, ZNC, MEV, NWS, MHR, ALO, CAL performed experiments and/or analyzed data. WG and DBL interpreted data, wrote and revised the paper. WG made the figures. DBL supervised overall design and study interpretation.

## Acknowledgements

Funding: Melanoma Research Alliance (DL and CAL), NIH R01GM101171, AACR-Bayer (17-80-44-LOMB), DoD awards CA170628, CA190267, OC140123, and NF170044 (DL), and the Rogel Cancer Center. Use of the Rogel Cancer Center shared resources is also gratefully acknowledged by DL and CAL (P30CA46592). SK was supported in part through an award from the Pablove Foundation. CAL was supported by a 2017 AACR NextGen Grant for Transformative Cancer Research (17-20-01-LYSS), an ACS Research Scholar Grant (RSG-18-186-01), and NIH award R01CA244931. ZNC was supported by NIH R01CA217141 and AACR-Bayer (19-80-44-NIKO). Metabolomics studies performed at the University of Michigan were supported by NIH grant DK097153. ALP is supported by the Highlands and Islands Enterprise (HMS9353763); MSS holds a fellowship (APP1106491) from the National Health and Medical Research Council (NHMRC). WG, AG, AM and LR were supported by NIH T32 awards (WG: GM007544, AG000114, HL007853 and AR007917; AG: AG000114 and GM113900; AM: AG000114; LR NL007517). NWS was supported by R01GM132261. ST was funded by American Diabetes Association postdoctoral fellowship #1-18-PDF-144. RAS and JSW are supported by the Australian National Health and Medical Research Council Fellowship program. RAS is also supported by a National Health and Medical Research Council of Australia Program Grant (APP1093017). Support from colleagues at Melanoma Institute Australia and the Royal Prince Alfred Hospital is also gratefully acknowledged. HPS holds an NHMRC MRFF Next Generation Clinical Researchers Program Practitioner Fellowship (APP1137127). The SBP Cancer Metabolism Core is supported by National Cancer Institute Cancer Center Support Grant P30 030199. Dr. Jeongsoon Park is acknowledged for technical contributions early in the project. Dr. Scott Pletcher (UM) is acknowledged for assistance with statistical analysis. Drs. Emily Bernstein (Icahn School of Medicine at Mount Sinai) and Kathleen Cho (UM) are acknowledged for generously providing cell lines. Drs. David Fisher (MGH Cancer Center/Harvard Medical School) and Kathryn Wellen (University of Pennsylvania) are gratefully acknowledged for helpful discussions. Olga Zagnitko is acknowledged for assistance with GC/MS measurements. Drs. Robert Weiss, Hening Lin, and Michael Deininger are acknowledged for discussion of unpublished data. GEO submission of RNA-seq data is in process.

