## Supplemental Figures S1-S7 for "The deacylase SIRT5 supports melanoma viability by regulating chromatin dynamics"

### Supplemental Figure 1

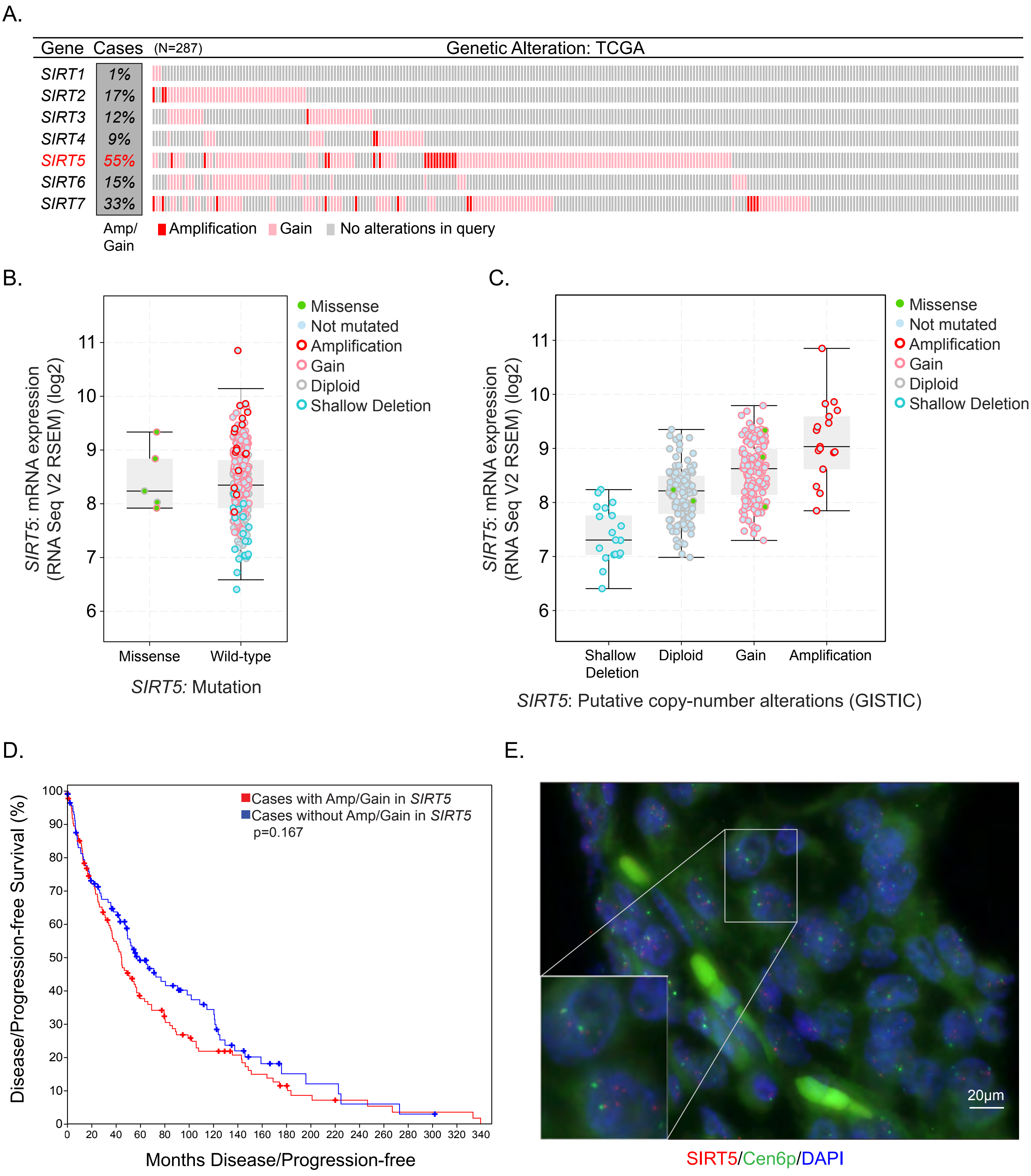

### Supplemental Figure 2

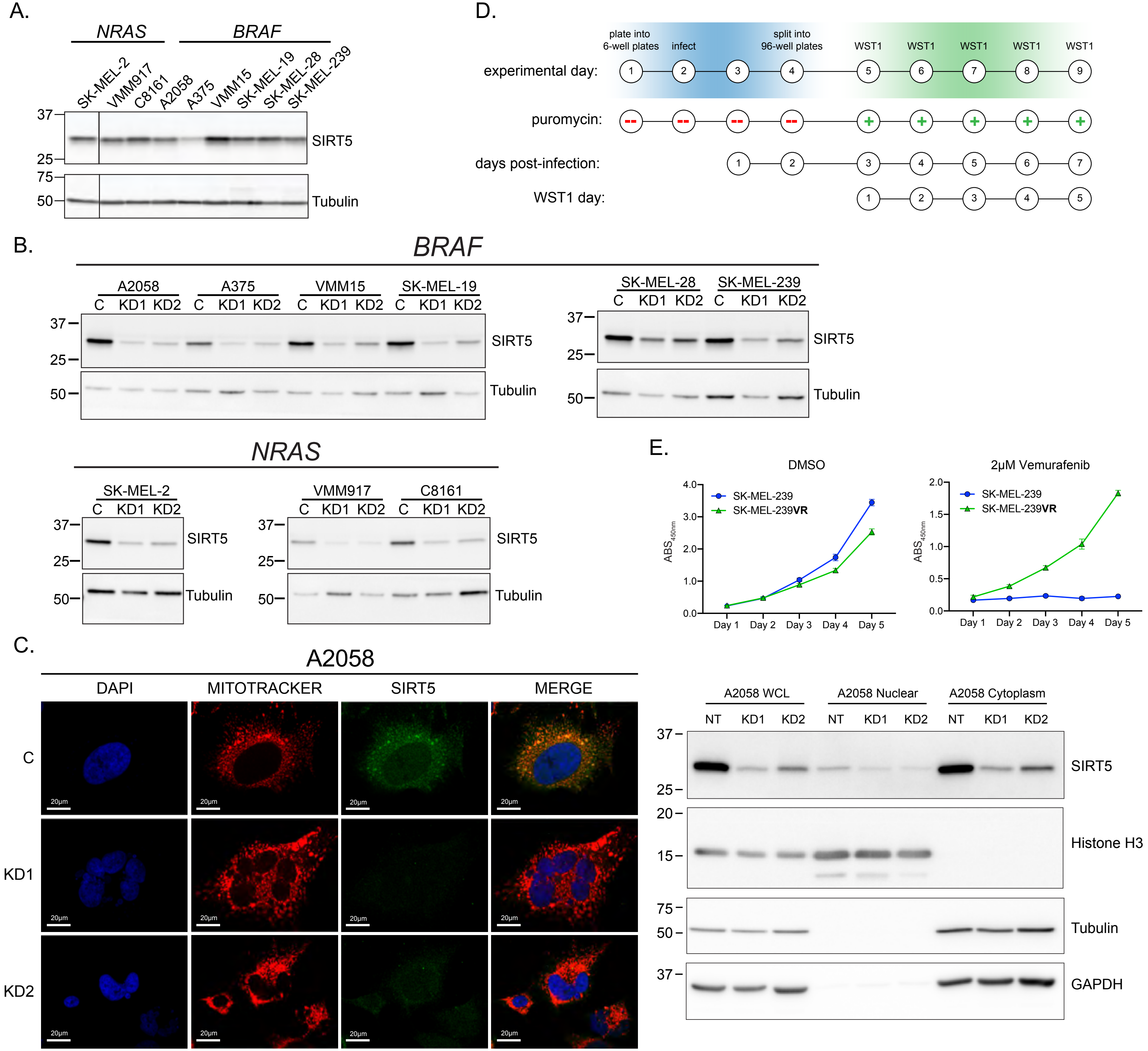

### Supplemental Figure 2

F.

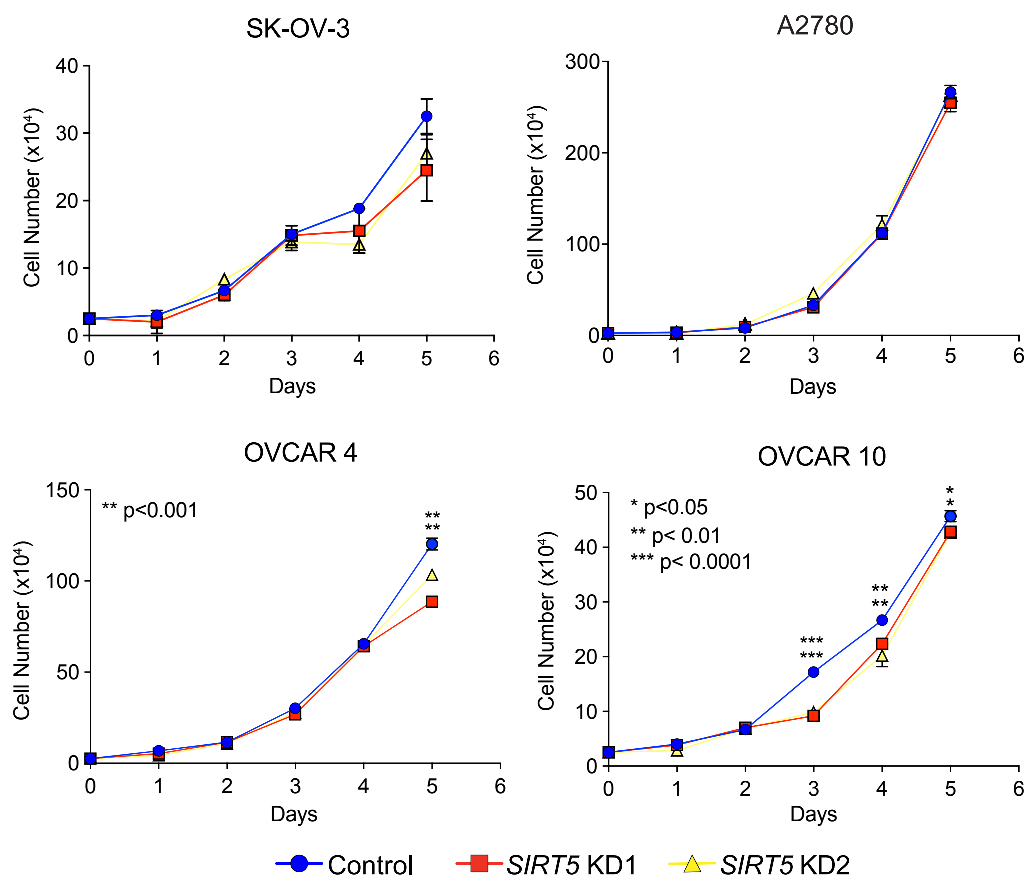


G.

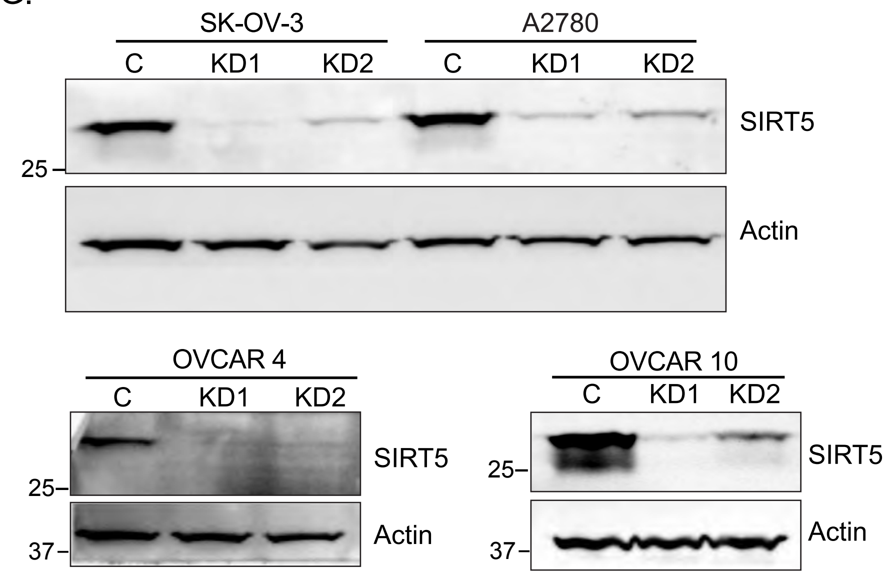

### Supplemental Figure 3

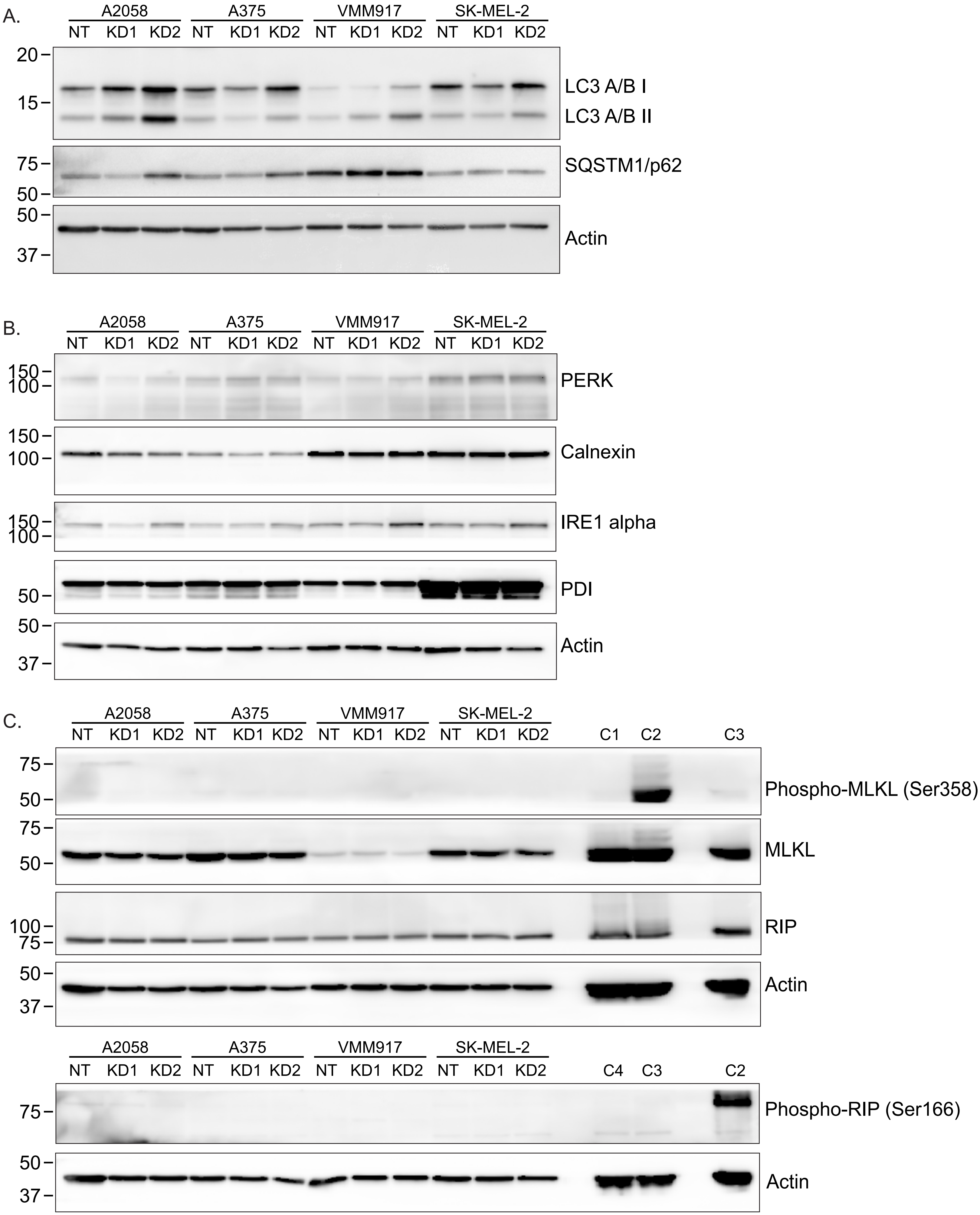

### Supplemental Figure 3

D.

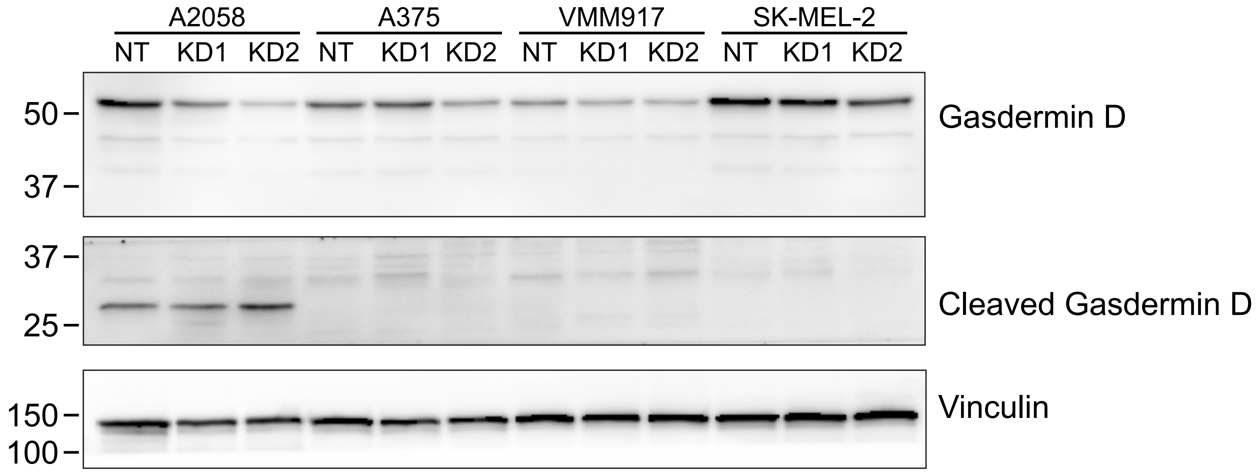


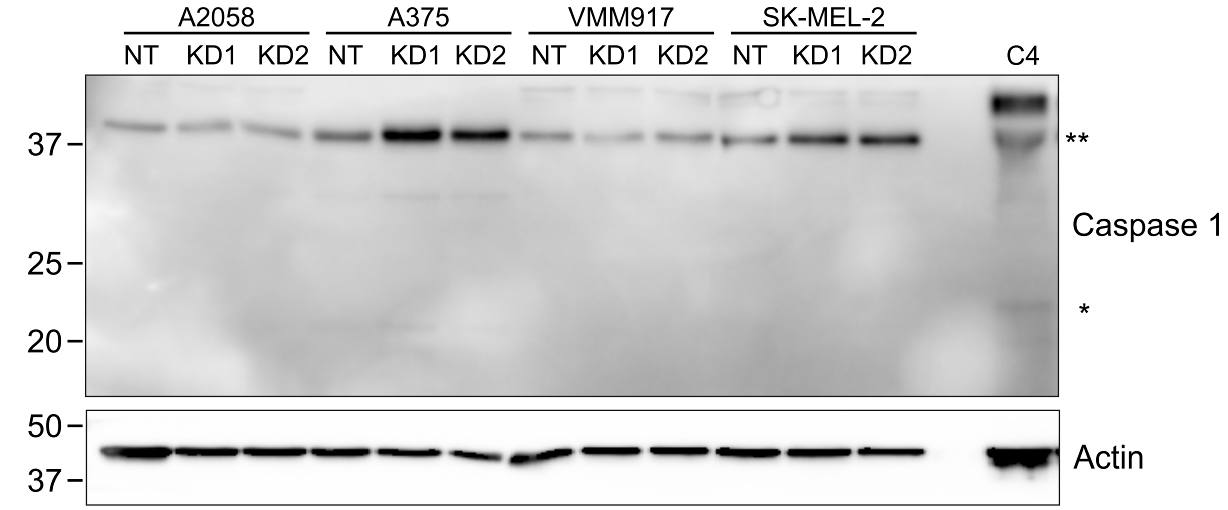

### Supplemental Figure 4

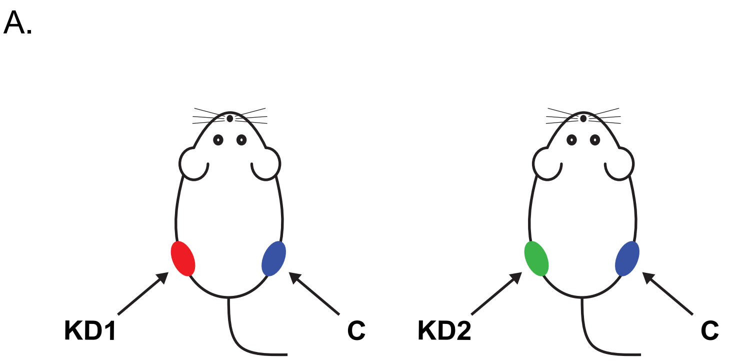


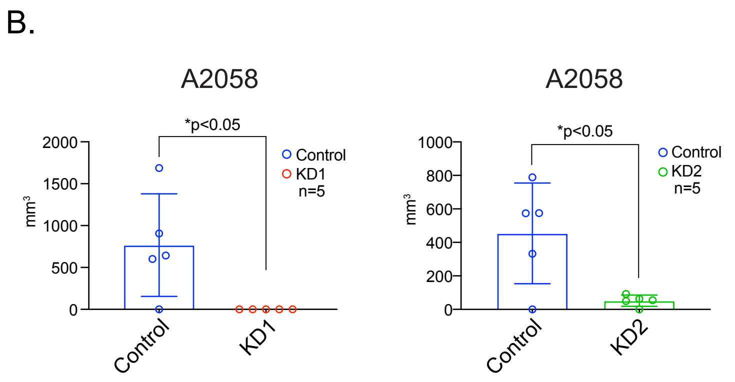

### Supplemental Figure 5

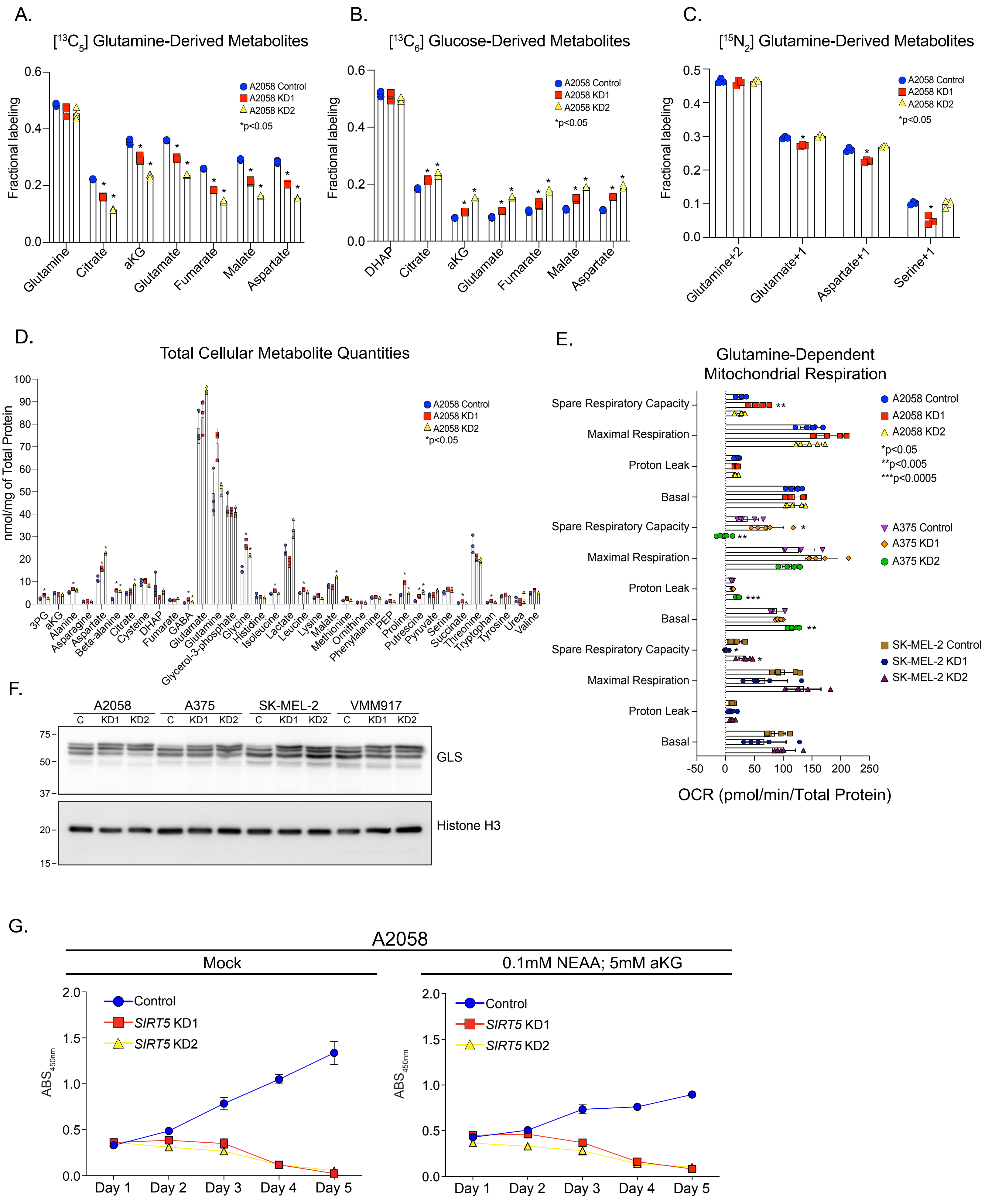

### Supplemental Figure 6

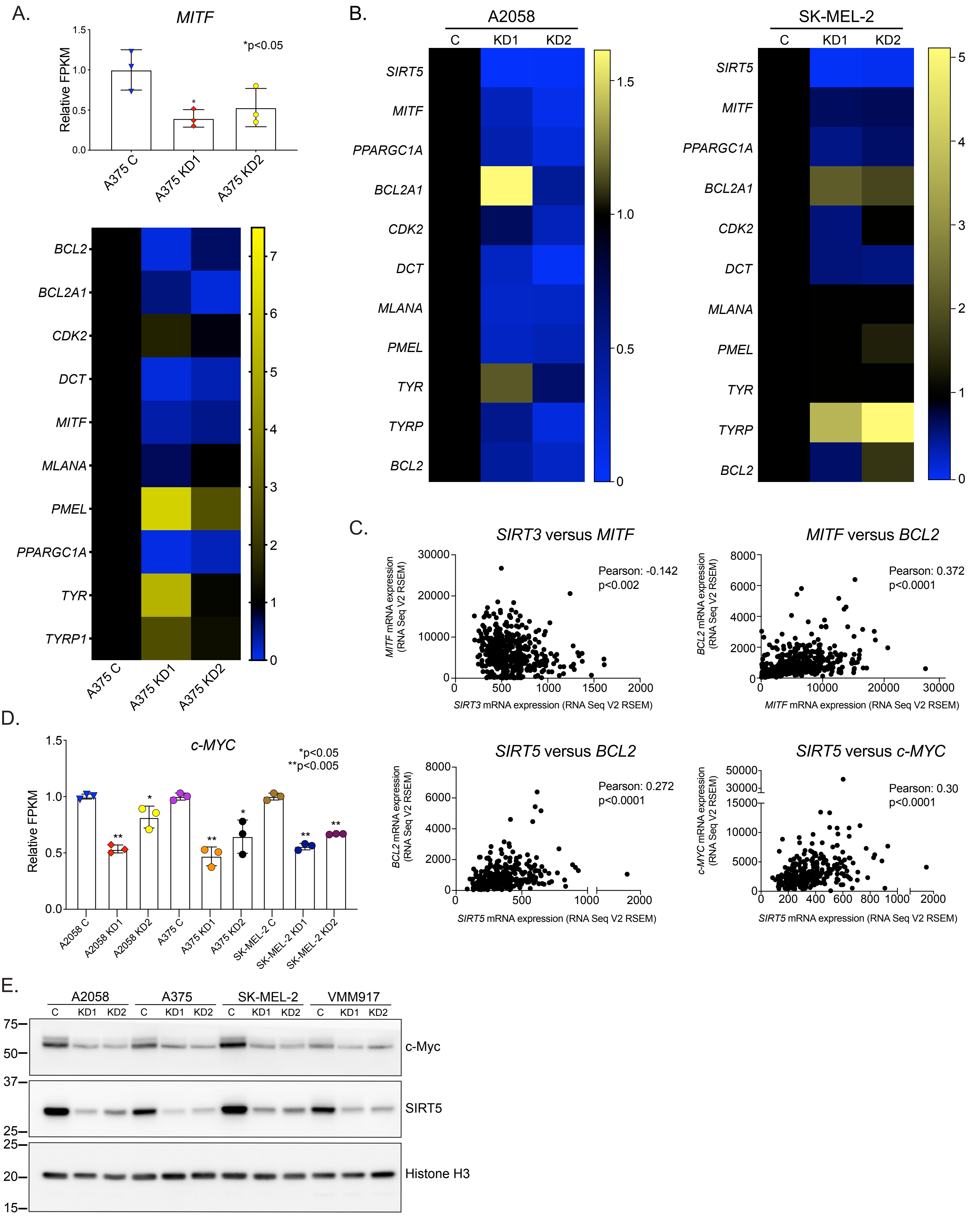

### Supplemental Figure 6

F.

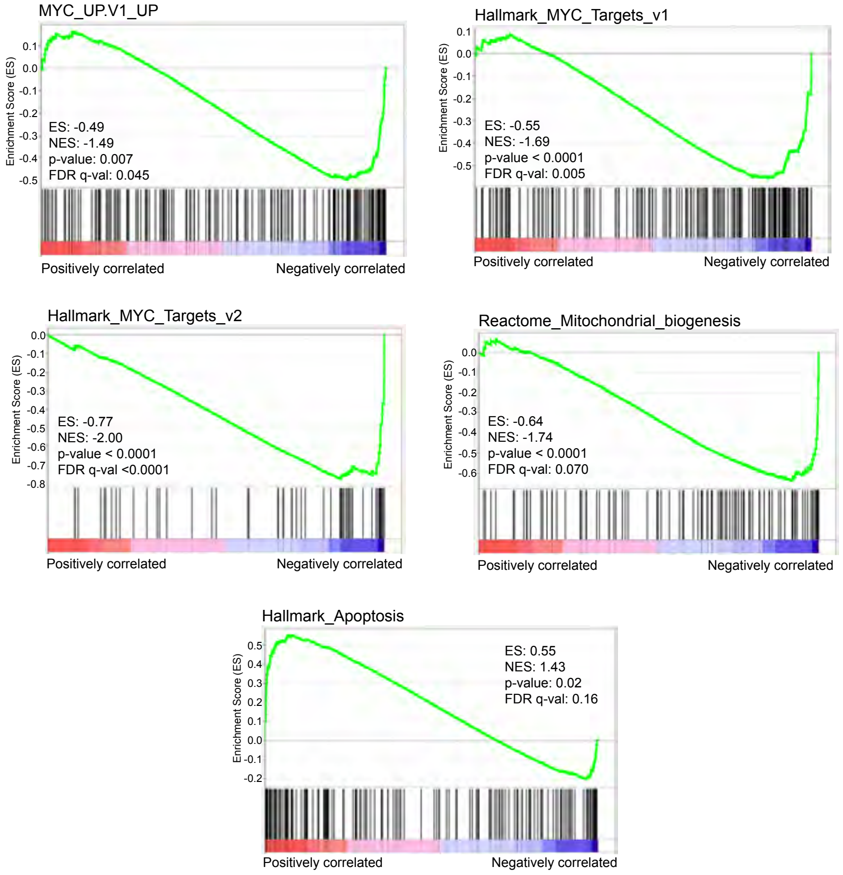


G.

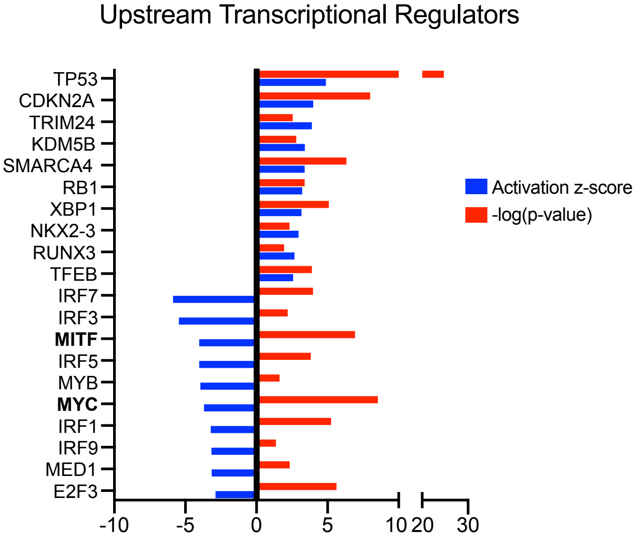


H.

| Ingenuity Canonical Pathways | -log(p-value) | Ratio | z-score |
|---|---|---|---|
| Molecular Mechanisms of Cancer | 10.5 | 0.395 | N/A |
| Senescence Pathway | 9.19 | 0.415 | 0.962 |
| Estrogen Receptor Signaling | 8.24 | 0.39 | 1.331 |
| PDGF Signaling | 6.44 | 0.5 | 2.16 |
| Cyclins and Cell Cycle Regulation | 6.36 | 0.506 | -3.053 |
| Pyridoxal 5'-phosphate Salvage Pathway | 6.14 | 0.53 | 0.686 |
| Sirtuin Signaling Pathway | 6.09 | 0.375 | 0.318 |
| Glioblastoma Multiforme Signaling | 6.01 | 0.418 | 0.397 |
| Aryl Hydrocarbon Receptor Signaling | 5.73 | 0.427 | -0.762 |
| Virus Entry via Endocytic Pathways | 5.67 | 0.461 | N/A |
| Integrin Signaling | 5.57 | 0.39 | 2.63 |
| IGF-1 Signaling | 5.38 | 0.452 | 2.333 |
| TGF-β Signaling | 5.28 | 0.458 | 0.16 |
| PPARα/RXRα Activation | 5.24 | 0.393 | -1.953 |
| Protein Ubiquitination Pathway | 5.15 | 0.366 | N/A |
| Salvage Pathways of Pyrimidine Ribonucleotides | 4.99 | 0.449 | 0.457 |
| Glioma Signaling | 4.97 | 0.436 | 1.257 |
| Cell Cycle: G1/S Checkpoint Regulation | 4.92 | 0.493 | 1.732 |
| Oncostatin M Signaling | 4.9 | 0.558 | 2.558 |
| Wnt/β-catenin Signaling | 4.83 | 0.393 | -0.662 |
| Insulin Receptor Signaling | 4.78 | 0.41 | 0.962 |
| Role of CHK Proteins in Cell Cycle Checkpoint Control | 4.74 | 0.509 | 1.528 |
| HER-2 Signaling in Breast Cancer | 4.66 | 0.381 | 0.117 |
| Hepatic Fibrosis Signaling Pathway | 4.65 | 0.341 | 1.008 |
| HIF1α Signaling | 4.55 | 0.376 | -0.114 |
| NRF2-mediated Oxidative Stress Response | 4.53 | 0.381 | 2.111 |
| Myc Mediated Apoptosis Signaling | 4.52 | 0.52 | 0.784 |
| Prolactin Signaling | 4.5 | 0.457 | 1.061 |
| Clathrin-mediated Endocytosis Signaling | 4.47 | 0.378 | N/A |
| Ovarian Cancer Signaling | 4.45 | 0.403 | 2.268 |
| 14-3-3-mediated Signaling | 4.4 | 0.409 | 0.316 |
| Rac Signaling | 4.38 | 0.413 | 4.025 |
| Pancreatic Adenocarcinoma Signaling | 4.35 | 0.422 | 0.169 |
| ErbB Signaling | 4.34 | 0.436 | 1.761 |
| ERK/MAPK Signaling | 4.26 | 0.371 | 1.588 |
| Renal Cell Carcinoma Signaling | 4.23 | 0.45 | 2.294 |
| Axonal Guidance Signaling | 4.21 | 0.324 | N/A |
| p53 Signaling | 4.21 | 0.429 | -0.898 |
| Huntington's Disease Signaling | 4.21 | 0.36 | -0.59 |
| Cell Cycle: G2/M DNA Damage Checkpoint Regulation | 4.19 | 0.51 | -1.279 |
| Estrogen-mediated S-phase Entry | 4.14 | 0.615 | -1.807 |
| PI3K/AKT Signaling | 4.12 | 0.375 | 0.277 |
| EGF Signaling | 4.1 | 0.491 | 2.502 |
| Endocannabinoid Cancer Inhibition Pathway | 4.04 | 0.392 | 0.816 |
| Autophagy | 4.04 | 0.475 | 0 |
| Non-Small Cell Lung Cancer Signaling | 3.97 | 0.452 | 1.633 |
| Tumor Microenvironment Pathway | 3.97 | 0.375 | 0.246 |
| Cell Cycle Regulation by BTG Family Proteins | 3.91 | 0.541 | -1.897 |
| Phagosome Maturation | 3.89 | 0.384 | N/A |
| Ferroptosis Signaling Pathway | 3.84 | 0.397 | 1 |

### Supplemental Figure 7

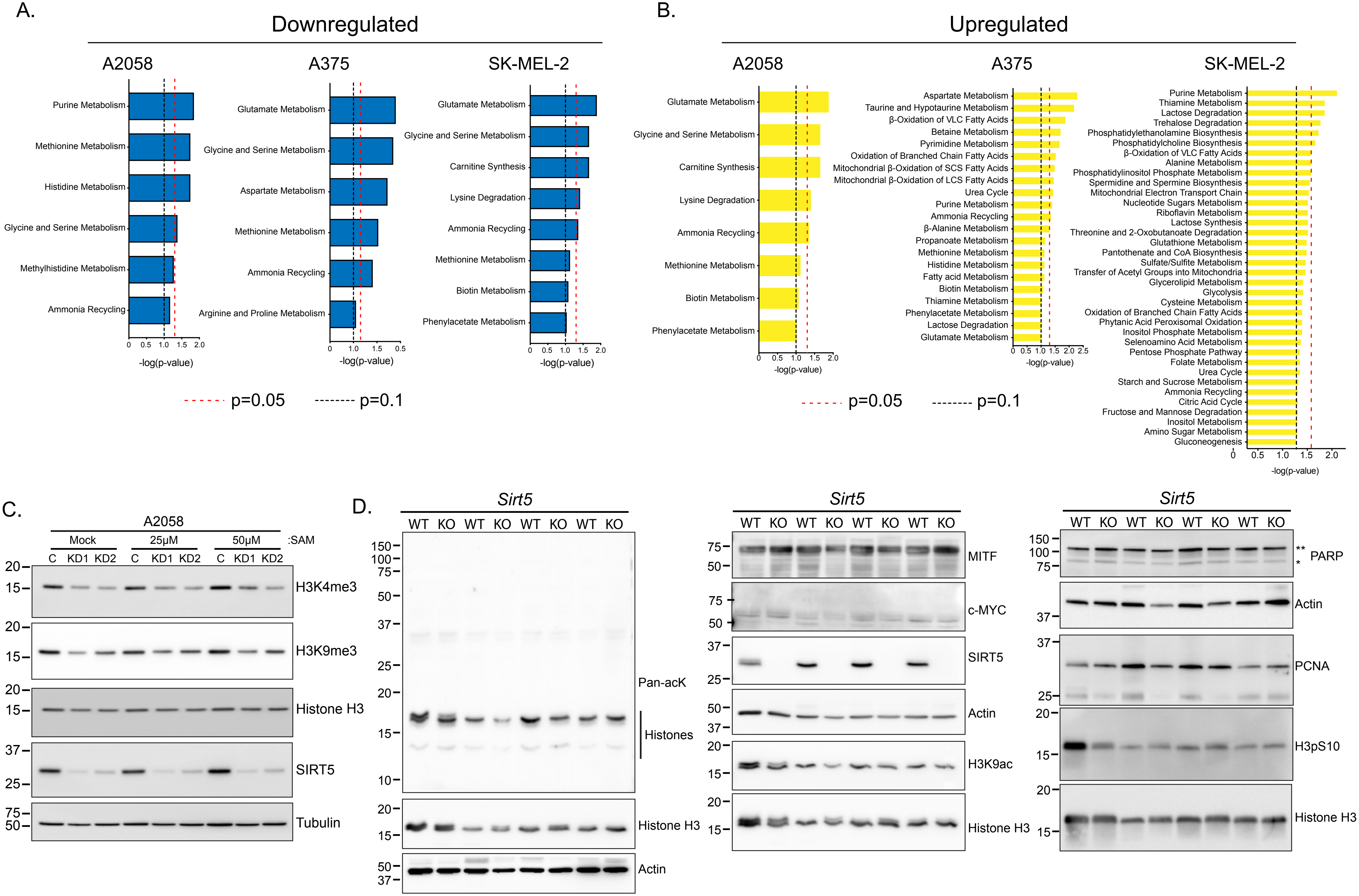
