## Supplemental Legends for "The deacylase SIRT5 supports melanoma viability by regulating chromatin dynamics"

**Figure S1. Sirtuin genetic alterations and disease-free survival in human melanoma patients.**

**A.** Gain and amplification of the *SIRT5* locus in melanoma. Other sirtuins are shown for comparison (n=287; data from TCGA, Provisional, analyzed on cBioPortal). Genomic amplification or gain of each sirtuin was analyzed using the query "*SIRT#*: AMP GAIN." **B.** The *SIRT5* sequence is WT in 98.3% (282/287) of melanoma samples (data from TCGA, Provisional, analyzed on cBioPortal). **C.** Increased *SIRT5* mRNA expression positively correlates with *SIRT5* gene gain or amplification in melanomas (data from TCGA, Provisional, analyzed on cBioPortal). *SIRT5* mRNA expression versus copy-number alterations (from GISTIC (1)) is plotted. **D.** Kaplan–Meier analysis of overall disease/progression-free survival in melanoma patients with or without gain or amplification of *SIRT5* (n=287; data from TCGA, Provisional, analyzed on cBioPortal). Survival was plotted using the query: "*SIRT5*: AMP GAIN." **E.** Representative FISH images of SIRT5 (red), centromere 6 (Cen6p) (green) and DAPI (blue) in a melanoma clinical sample. Scale bar=20μm.

**Figure S2. SIRT5 is expressed in human melanoma and ovarian cancer cell lines.**

**A.** SIRT5 protein expression in human melanoma cell lines used in this study. Lanes were run on the same gel but are noncontiguous. **B.** Representative immunoblot analysis showing SIRT5 protein levels in melanoma cells 96 hrs. post-transduction with a non-targeting shRNA (C) or one of two SIRT5 shRNAs (KD1 or KD2). **C.** Confocal immunofluorescence and immunoblot analysis of SIRT5 protein in A2058 cells showing SIRT5 depletion in mitochondria, cytosol, and nucleus following SIRT5 KD. DAPI (blue), Mitotracker (red), SIRT5 (green). **D.** Cell viability assay schematic. **E.** SK-MEL-239**VR** cells are resistant to vemurafenib treatment compared to vemurafenib-sensitive SK-MEL-239 cells. Equivalent cell numbers were plated into 96-well plates in the presence of vehicle (DMSO) or 2μM vemurafenib (n=6/timepoint). Cell mass was determined at the indicated timepoints via WST-1 assay, with absorbance measured at 450nm. **F.** Ovarian cancer cell lines, as indicated, were infected with a non-targeting shRNA (control) or one of two SIRT5 shRNAs (KD1 or KD2). Equivalent cell numbers were then plated into 96-well plates in the presence of puromycin in triplicates into 12-well plates. Cell number in each well were assessed every 24 hrs. after plating by manual counting using a hemocytometer. Error bars represent standard deviation. Significance calculated using unpaired Student's t-test. **G.** Representative immunoblot analysis showing SIRT5 protein levels in indicated ovarian cell lines post-transduction with a non-targeting shRNA (C) or one of two SIRT5 shRNAs (KD1 or KD2).

**Figure S3. Analysis of markers non-apoptotic cell death upon SIRT5 KD.**

Markers for autophagic- (**A.**), ER-stress- (**B.**), necroptotic- (**C.**) and pyroptotic- (**D.**) induced cell death were assessed in the indicated melanoma cell lines by immunoblot. Control lysates: C1, untreated HT29 cells; C2, HT29 cells treated with Z-VAD (20 µM for 30 min), followed by addition of cycloheximide (10 µg/ml) and TNFα (20 ng/ml) overnight; C3: THP1 cells treated with TPA (80nM overnight) and LPS (1 µg/ml for 7 hr); C4, untreated THP1 cells. **, full-length caspase-1; *, caspase-1 cleavage product. All control lysates were provided by Cell Signaling Technology upon request. No specific signal was observed for p-PERK (not shown).

**Figure S4. SIRT5 depletion reduces tumor formation in mouse xenografts.**

**A.** Non-targeting (C) or SIRT5 KD (KD1 or KD2) cells were injected into the right or left flanks, respectively, of immunocompromised mice as indicated. **B.** Plotted are averages (n=5) of tumor volume measurements in $mm^3$ taken at day 28. Each dot represents individual endpoint tumor size in each mouse. Error bars represent SD; $p<0.05$, paired two-tailed t-test.

**Figure S5. Metabolic perturbations in melanoma cells upon SIRT5 loss.**

Fractionally-labeled [$^{13}C_5$]-glutamine-derived metabolite (**A.**) and [$^{13}C_6$]-glucose-derived metabolite (**B.**) levels upon SIRT5 depletion in A2058 cells. **C.** [$^{15}N_2$]-glutamine-derived metabolite levels upon SIRT5 depletion in A2058 cells quantified and plotted as in (A.). n=3 for each sample. Error bars represent standard deviation. **D.** Representative unlabeled, total cellular metabolite quantities in control and SIRT5 knockdown samples. Plotted are average metabolite levels in nmol metabolite/mg of total protein. aKG; alpha-ketoglutarate, 3PG; 3-phosphoglyceric acid, DHAP; dihydroxyacetone phosphate, GABA; gamma-aminobutyric acid, PEP; phosphoenolpyruvate. n=3 for each sample. Error bars represent standard deviation. **E.** Glutamine-dependent mitochondrial respiration in A2058, A375 and SK-MEL-2 cells (n=5) is unchanged upon SIRT5 ablation. Error bars represent standard deviation. **F.** Glutaminase (GLS) immunoblot in melanoma cell lines, as indicated, 96 hrs. post-transduction with shRNAs targeting *SIRT5* (KD1 or KD2) compared to a non-targeting control (C). Histone H3 serves as the loading control. **G.** A2058 cells were infected with a non-targeting shRNA (control) or one of two SIRT5 shRNAs (KD1 or KD2). Equivalent cell numbers were then plated 48 hrs. post-transduction into 96-well plates in the absence (left panel) or presence (right panel) of 0.1mM non-essential amino acids (NEAA) and 5mM aKG. Cell mass was determined at the indicated timepoints via WST-1 assay, with absorbance measured at 450nm. Average results (n=6/timepoint) are graphed. Error bars represent standard deviation. Significance calculated using unpaired Student's t-test.

**Figure S6. Altered MITF and c-MYC pathway signaling in SIRT5-depleted melanoma.**

**A.** Reduced expression of *MITF* (bar graph, upper panel) and *MITF* target gene transcripts upon SIRT5 KD in A375 cells (heatmap, lower panel). Scale bars adjacent to heat maps indicate linear fold change (control (C) set to 1). Error bars represent standard deviation. Significance calculated using unpaired Student's t-test. **B.** Confirmation of attenuated mRNA expression of *SIRT5*, *MITF* and *MITF* responsive genes by qRT-PCR in A2058 and SK-MEL-2 cells (n=3).

Scale bars adjacent to heat maps indicate linear fold change (control (C) set to 1). **C.** Pairwise correlations of *SIRT3*, *SIRT5*, *MITF, c-MYC* and the *MITF* target, *BCL2* in melanoma clinical samples (data from TCGA, analyzed on cBioPortal; see Figure 1A). **D.** Relative FPKM values in A2058, A375 and SK-MEL-2 cells demonstrate reduced *c-MYC* mRNA expression upon SIRT5 KD. Error bars represent standard deviation. Significance calculated using unpaired Student's t-test. **E.** Immunoblot demonstrating loss of c-MYC protein expression 96 hrs. post-transduction with shRNAs targeting *SIRT5* (KD1 or KD2) compared to a non-targeting control (C) in 4 cutaneous *BRAF*- (A2058 and A375) or *NRAS*- (SK-MEL-2 and VMM917) mutant melanoma cell lines. **F.** GSEA of aggregate *SIRT5* knockdown samples compared to *SIRT5*-proficient controls reveals a negative enrichment of *c-MYC*, c-MYC target genes, and mitochondrial biogenesis pathways. Positive enrichment of genes involved in apoptosis was observed in the same aggregate *SIRT5* knockdown data set. **G.** The top 20 transcriptional regulators, sorted by activation z-score, predicted to be either activated or inhibited upon SIRT5 loss when comparing aggregate *SIRT5* knockdown samples to controls. The activation z-score and significance are graphed. All graphed regulators meet significance thresholds set by Ingenuity Pathway Analysis ($p<0.05$ and absolute z-score>2). **H.** Top 50 pathways significantly enriched by differentially expressed genes in SIRT5 knockdown versus control samples (aggregate of all cell lines analyzed), using Ingenuity Pathway Analysis (IPA). Ingenuity Canonical Pathway, pathway name; -log(p-value), significance of pathway; Ratio, number of genes represented in the experimental list compared to the total number of genes mapped to that pathway; z-score, IPA prediction on whether the pathway is up or downregulated based on the differentially expressed genes mapped. IPA does not generate z-scores on all pathways; these are labelled N/A.

**Figure S7. Metabolic pathway analysis reveals altered metabolism following SIRT5 loss.** MetaboAnalyst pathway analysis of downregulated (**A.**) or upregulated (**B.**) metabolites altered in both SIRT5 KD1 and KD2 samples of the indicated melanoma cell lines. Dashed lines represent a p-value cutoff of $p<0.05$ or $p<0.1$, as indicated. **C.** Incubation of SIRT5-depleted A2058 cells with 25μM or 50μM SAM in culture for 24hrs. does not restore H3K4me3 or H3K9me3 levels. **D.** Immunoblot reveals histone acetylation, MITF and c-MYC protein expression are restored in tumors arising in SIRT5-deficient $\textit{Braf}^{CA}$;$\textit{Pten}^{fl/fl}$;*Tyr::CreER* mice (see Figure 4C). Markers for cell death (PARP cleavage) and mitosis (PCNA and histone H3 phosopho-S10 (H3pS10)) are consistent in tumors arising in SIRT5-deficient $\textit{Braf}^{CA}$;$\textit{Pten}^{fl/fl}$;*Tyr::CreER* mice. **, full-length PARP; *, PARP cleavage product.

### Supplemental Tables

**Table S1.** Copy number aberrations in SIRT1-7 in benign and dysplastic nevi. LOH, Loss of heterozygosity; HD, Homozygous deletion.

**Table S2.** Cutaneous and uveal cell lines used in this study. Name, source and catalog number, where available, are listed. Mutations in major melanoma-relevant genes, if known, are indicated. Sex and age of patient at the time of cell line derivation, if known, are listed.

**Table S3.** FPKM values from RNAseq analysis for each sample (control (C) or SIRT5 knockdown (KD1 or KD2)), as indicated, is listed for each gene. FPKM, fragments per kilobase of transcript per million mapped reads. DEseq analysis, comparing changes in gene expression in control to those common to both KDs is listed for A2058, A375 and SK-MEL-2 cell lines. PADJ, p-value adjusted for multiple comparisons.

**Table S4.** Flux data generated from genome-scale metabolic modeling. The 20 significant reactions and raw data are listed.

**Table S5.** Raw ion currents from LC-MS/MS metabolite profiling in control (C) and SIRT5 knockdown (KD1 or KD2) samples. QqQ, triple quadrupole; Compound Method, metabolite detected; Transition, Parent m/z > Fragment m/z; m/z: mass to charge ratio.

**Table S6.** Human and mouse shRNA sequences used to deplete SIRT5 are listed along with The RNAi Consortium (TRC) clone numbers where indicated. Human KD3 and KD4 shRNAs were designed based on published CRISPR databases (see methods). CRISPR sequences targeting human *SIRT5* are listed.

**Table S7.** Antibodies used in this study. Antibody name, source, catalog number and Research Resource Identifiers (RRIDs) are indicated.
