## Supplementary material for "The deacylase SIRT5 supports melanoma viability by regulating chromatin dynamics": Table S1

| Sample ID | SIRT1 | SIRT2 | SIRT3 | SIRT4 | SIRT5 | SIRT6 | SIRT7 | Dermoscopic Pattern | Histopath diagnosis | Histopathological Architecture | Complete Histopath diagnosis |
| --- | --- | --- | --- | --- | --- | --- | --- | --- | --- | --- | --- |
| 52JH-40 |  |  |  |  |  |  |  | Globular | Benign | Compound | benign compound nevus, junctional component minimal |
| 772DW-75 |  |  |  |  |  |  |  | Reticular | Benign | Compound | compound nevus |
| 07LW-152 |  |  |  |  |  |  |  | Globular | Dysplastic | Compound | compound dysplastic nevus |
| 07LW-155 |  |  |  | LOH^a^ |  | LOH | LOH | Reticular | Dysplastic | Junctional | junctional dysplastic nevus |
| 158RP-4 |  |  |  |  |  |  |  | Globular | Benign | Intradermal | benign intradermal nevus extending to base of specimen |
| 111CM-3 |  |  |  |  |  |  |  | Globular | Benign | Compound | benign compound nevus with periappendageal accentuation focally involving base of shave |
| 130MV-160 | LOH | HD^b^ |  | HD |  | LOH |  | Reticular | Benign | Compound | benign lentiginous compound nevus of superficial congenital type |
| 903JA-33 |  |  |  |  |  |  |  | Reticular | Dysplastic | Junctional | junctional dysplastic nevus |
| 111CM-21 |  |  |  |  |  |  |  | Globular | Benign | Intradermal | benign intradermal nevus focally extending to edge of specimen |
| 28RH-135 |  |  |  |  |  |  |  | Globular | Benign | Intradermal | intradermal nevus |
| 61JB-133 |  |  |  |  |  |  |  | Globular | Benign | Intradermal | benign intradermal nevus with congenital features |
| 736TP-3 |  |  |  |  |  |  |  | Globular | Dysplastic | Compound | compound dysplastic nevus with mild atypia |
| 1346KJ-30 |  |  |  |  |  |  |  | Globular | Benign | Compound | compound nevus with congenital features |
| 673PS-1 |  |  | LOH |  |  |  |  | Reticular | Benign | Junctional | junctional lentiginous nevus |
| 736TP-13 | LOH | LOH |  | LOH | LOH | LOH | LOH | Reticular | Benign | Junctional | junctional lentiginous nevus |
| 28RH-110 |  |  |  |  |  |  |  | Reticular | Dysplastic | Compound | compound dysplastic nevus with mild atypia |
| 155IN-30 |  |  |  |  |  |  |  | Reticular | Benign | Compound | compound nevus |
| 158RP-78 |  |  | LOH |  | LOH |  |  | Reticular | Benign | Junctional | junctional nevus extending to peripheral edge of specimen |
| 510AN-2 |  |  |  |  |  |  |  | Globular | Benign | Intradermal | benign intradermal nevus |
| 39RB-6 |  |  |  |  |  |  |  | Globular | Benign | Intradermal | intradermal nevus |
| 61JB-115 |  |  |  |  |  |  |  | Globular | Benign | Compound | benign compound nevus with congenital features, minimal epidermal component |
| 1062DM-76 |  |  |  |  |  |  |  | Non-specific | Dysplastic | Compound | dysplastic compound nevus with mild atypia |
| 822MT-10 |  |  |  |  |  |  |  | Non-specific | Dysplastic | Compound | compound dysplastic nevus with mild atypia |
| 1078JG-100 |  | LOH |  |  | LOH | LOH | LOH | Non-specific | Dysplastic | Compound | compound dysplastic nevus with mild atypia |
| 695MB-59 |  |  |  |  |  |  |  | Reticular | Dysplastic | Compound | compound dysplastic nevus |
| 1346KJ-7 |  |  |  |  |  |  |  | Reticular | Benign | Compound | compound nevus |
| 774AS-17 |  |  |  |  |  |  |  | Reticular | Benign | Junctional | junctional lentiginous nevus |
| 673PS-2 | LOH |  |  |  |  |  |  | Reticular | Dysplastic | Junctional | junctional dysplastic nevus |
| 822MT-RLA | LOH |  |  | LOH |  | LOH | LOH | Non-specific | Dysplastic | Junctional | junctional dysplastic nevus with mild atypia |
| 510AN-23 |  | LOH |  | LOH |  | LOH |  | Reticular | Dysplastic | Compound | compound dysplastic nevus with mild atypia |

Table S1

^a^LOH, Loss of heterozygosity

^b^HD, Homozygous deletion
