## Supplementary material for "The deacylase SIRT5 supports melanoma viability by regulating chromatin dynamics": Table S2

| **Cell Line** | **Source/Catalog Number** | **BRAF** | **NRAS** | **Age** | **Sex** |
| --- | --- | --- | --- | --- | --- |
| **Cutaneous** |  |  |  |  |  |
| A375 | ATCC CRL-1619 | V600E | WT | 54 | F |
| A2058 | ATCC CRL-11147 | V600E | WT | 43 | M |
| SK-MEL-28 | ATCC HTB-72 | V600E | WT | 51 | M |
| VMM15 | ATCC CRL-3227 | V600E | WT | 39 | M |
| SK-MEL-19 | -- | V600E | WT |  |  |
| SK-MEL-239 | -- | V600E |  |  |  |
| SK-MEL-239**VR** |  | V600E |  |  |  |
| SK-MEL-2 | ATCC HTB-68 | WT | Q61R | 60 | M |
| VMM917 | ATCC CRL-3232 | WT | Q61L, Q61R | 57 | M |
| C8161 | -- | WT | Q61R |  |  |
| **Uveal** |  | **GNA11** | **GNAQ** |  |  |
| MP-38 | ATCC CRL-3296 |  |  |  | M |
| MP-41 | ATCC CRL-3297 | c.626a A>T |  | 49 | F |
| MP-46 | ATCC CRL-3298 |  | c.626a A>T | 69 | M |
| 92-1 | Sigma 13012458 |  |  | 79 | F |

Table S2
