## Supplementary material for "The deacylase SIRT5 supports melanoma viability by regulating chromatin dynamics": Table S6

| Name | Sequence | Clone Name |
| --- | --- | --- |
| Human *SIRT5* shRNA |  |  |
| KD1 | 5’-AAATCTGGTTTCGTGTGGACG-3’ | TRCN0000018546 |
| KD2 | 5’ TTTCTGCACTAACACCAGCTC-3’ | TRCN0000018547 |
| KD3 | 5’-TCTCACACTCGGCTATGGCG-3’ | N/A |
| KD4 | 5’-GGCTGCTGGGTACACCACAG-3’, | N/A |
| KD5 | 5’-TTTAGATTGTTCAGTACTCAG-3’ | TRCN0000232662 |
| Mouse *Sirt5* shRNA |  |  |
| KD1 | 5’-ATCACGTAACAGATTGTCTGC-3’ | TRCN0000092833 |
| KD2 | 5’-TTTCGTCTACAACACAACTGG-3’ | TRCN0000092834 |
| KD3 | 5’-AAATGAAACCTGAATCTGTCG-3’ | TRCN0000092835 |
| KD4 | 5’-ATCGGACTCCTATAGTTCTCG-3’ | TRCN0000092836 |
| KD5 | 5’-AAAGTCTGCCATATTTGAACT-3’ | TRCN0000092837 |
| Human *SIRT5* CRISPR Guides |  |  |
| G1 | 5’-TCTCACACTCGGCTATGGCG-3’ | N/A |
| G2 | 5’-GGCTGCTGGGTACACCACAG-3’ | N/A |
| G3 | 5’-GGCTGCCAGGGGCGTGCCAG-3’ | N/A |
| G4 | 5’-GTTCCGACCTTCAGAGGAGC-3’ | N/A |
