## Supplementary material for "The deacylase SIRT5 supports melanoma viability by regulating chromatin dynamics": Table S7

| Antibodies | Source | Identifier |
| --- | --- | --- |
| Mouse monoclonal anti-alpha-Tubulin | Santa Cruz Biotechnology | Cat# sc-23948; RRID:AB_628410 |
| Rabbit monoclonal anti-MITF | Cell Signaling Technology | Cat# 12590, RRID:AB_2616024 |
| Rabbit monoclonal anti-Glutaminase | Abcam | Cat# ab156876, RRID:AB_2721038 |
| Rabbit monoclonal anti-SIRT5 | Cell Signaling Technology | Cat# 8779, N/A |
| Rabbit polyclonal anti-SIRT5 | Sigma-Aldrich | Cat# HPA022002, RRID:AB_1856913 |
| Rabbit monoclonal anti-HA | Cell Signaling Technology | Cat# 3724, RRID:AB_1549585 |
| Mouse monoclonal anti-SIRT5 (biotinylated) | Thermo Fisher Scientific | Cat# 730092, RRID:AB_2532891 |
| Rabbit monoclonal anti-Cleaved Caspase-3 (Asp175) (5A1E) | Cell Signaling Technology | Cat# 9664, RRID:AB_2070042 |
| Rabbit polyclonal anti-Acetylated Lysine | Cell Signaling Technology | Cat# 9441, RRID:AB_331805 |
| Mouse monoclonal anti-Histone H3 | Cell Signaling Technology | Cat# 14269, RRID:AB_2756816 |
| Mouse monoclonal anti-beta-Actin | Sigma-Aldrich | Cat# A5441, RRID:AB_476744 |
| Rabbit polyclonal anti-Histone H3 | Abcam | Cat# ab1791, RRID:AB_302613 |
| Rabbit polyclonal anti-Histone H3K9me3 | Abcam | Cat# ab8898, RRID:AB_306848 |
| Rabbit polyclonal anti-Histone H3K4me3 | Abcam | Cat# ab8580, RRID:AB_306649 |
| Rabbit polyclonal anti-Histone H3K9ac | Abcam | Cat# ab4441, RRID:AB_2118292 |
| Rabbit monoclonal anti-Histone H4K16ac | Cell Signaling Technology | Cat# 13534, RRID:AB_2687581 |
| Rabbit polyclonal anti-Histone H4 | Cell Signaling Technology | Cat# 2592, RRID:AB_2118614 |
| Rabbit monoclonal anti-c-Myc | Cell Signaling Technology | Cat# 5605, RRID:AB_1903938 |
| Rabbit polyclonal anti-MTHFD1L | Proteintech | Cat# 16113-1-AP, RRID:AB_2250974 |
| Alexa Fluor 488 goat anti-Rabbit IgG (H+L) | Thermo Fisher Scientific | Cat# A-11034, RRID:AB_2576217 |
| Normal Rabbit IgG | Cell Signaling Technology | Cat# 2729, RRID:AB_1031062 |
| Rabbit (DA1E) mAb IgG XP® Isotype Control (CUT&RUN) | Cell Signaling Technology | Cat# 3900, RRID:AB_1550038 |
| Rabbit Anti-Histone H3, acetyl (Lys9) Monoclonal Antibody, Unconjugated, Clone C5B11 (CUT&RUN) | Cell Signaling Technology | Cat# 9649, RRID:AB_823528 |
| Peroxidase AffiniPure Goat Anti-Mouse IgG (H+L) | Jackson ImmunoResearch Labs | Cat# 115-035-166, RRID:AB_2338511 |
| Peroxidase AffiniPure Goat Anti-Rabbit IgG (H+L) | Jackson ImmunoResearch Labs | Cat# 111-035-045, RRID:AB_2337938 |
| Rabbit monoclonal anti-MLKL | Cell Signaling Technology | Cat# 14993, RRID:AB_2721822 |
| Rabbit monoclonal anti-phospho-MLKL (Ser358) | Cell Signaling Technology | Cat# 91689, RRID:AB_2732034 |
| Rabbit monoclonal anti-RIP | Cell Signaling Technology | Cat# 3493, RRID:AB_2305314 |
| Rabbit monoclonal anti-phospho-RIP (Ser166) | Cell Signaling Technology | Cat# 65746, RRID:AB_2799693 |
| Rabbit monoclonal anti-PERK | Cell Signaling Technology | Cat# 5683, RRID:AB_10841299 |
| Rabbit monoclonal anti-Calnexin | Cell Signaling Technology | Cat# 2679, RRID:AB_2228381 |
| Rabbit monoclonal anti-IRE1 alpha | Cell Signaling Technology | Cat# 3294, RRID:AB_823545 |
| Rabbit monoclonal anti-PDI | Cell Signaling Technology | Cat# 3501, RRID:AB_2156433 |
| Rabbit monoclonal anti-Gasdermin D | Cell Signaling Technology | Cat# 97558, RRID:AB_2864253 |
| Rabbit monoclonal anti-Cleaved Gasdermin D (Asp275) | Cell Signaling Technology | Cat# 36425, RRID:AB_2799099 |
| Rabbit monoclonal anti-Caspase-1 | Cell Signaling Technology | Cat# 3866, RRID:AB_2069051 |
| Rabbit polyclonal anti-LC3B | Novus Biologicals | Cat# NB600-1384, RRID:AB_669581 |
| Mouse monoclonal anti-Vinculin | Santa Cruz Biotechnology | Cat# sc-73614, RRID:AB_1131294 |
| Rabbit polyclonal anti-SQSTM1/p62 | Cell Signaling Technology | Cat# 5114, RRID:AB_10624872 |
| Rabbit polyclonal anti-PARP | Cell Signaling Technology | Cat# 9542, RRID:AB_2160739 |
| Mouse monoclonal anti-PCNA | Abcam | Cat# ab29, RRID:AB_303394 |
| Rabbit monoclonal anti-phospho-Histone H3 (Ser10), Clone D2C8 | Cell Signaling Technology | Cat# 3377, RRID:AB_1549592 |
